# A Transcriptomic and Proteomic Atlas of Obesity and Type 2 Diabetes in Cynomolgus Monkeys

**DOI:** 10.1101/2021.12.10.472179

**Authors:** Xianglong Zhang, Ying Lei, Oliver Homann, Marina Stolina, Songli Wang, Murielle M. Véniant, Liangbiao George Hu, Yi-Hsiang Hsu

**Author notes:** Correspondence (Y.-H.H.), (L.G.H.), (M.M.V.). These authors contributed equally (Xianglong Zhang, Ying Lei).

## Abstract

Obesity and type 2 diabetes (T2D) remain major global healthcare challenges and developing therapeutics necessitate using nonhuman primate models. Here, we present transcriptomic and proteomic analyses of all the major organs of cynomolgus monkeys with spontaneous obesity or T2D in comparison to healthy controls. Molecular changes occur predominantly in the adipose tissues of individuals with obesity, while extensive expression perturbations among T2D individuals are observed in many tissues, such as the liver, kidney, brain, and heart. Immune response-related pathways are upregulated in obesity and T2D, whereas metabolism and mitochondrial pathways are downregulated. Incorporating human single-cell RNA sequencing findings corroborates the role of macrophages and monocytes in obesity. Moreover, we highlight some potential therapeutic targets including *SLC2A1* and *PCSK1* in obesity as well as *SLC30A8* and *SLC2A2* in T2D. Our findings provide insights into tissue-specific molecular foundations of obesity and T2D and reveal the mechanistic links between these two metabolic disorders.

## INTRODUCTION

Obesity and one of its associated severe comorbidities—type 2 diabetes (T2D)—pose a major health threat globally and have reached epidemic proportions over the past decades. Heritability estimates are around 40–70% for body mass index (BMI, a commonly used proxy phenotype for obesity) ^1^ and 30–70% for T2D ^2^. Genome-wide association studies (GWAS) have identified more than 300 loci associated with obesity and more than 3000 with T2D according to the GWAS catalog ^2^. However, the molecular mechanisms of how these loci regulate gene expression and impact protein function remain poorly understood.

Previous studies have investigated the transcriptomes or proteomes of obese patients in tissues including blood ^3,4^, adipose tissues ^5-8^, liver ^9^, muscle ^8,10,11^, breast ^12^, granulosa cells ^13^, and sperm^14^. However, these studies have yielded inconclusive and sometimes conflicting results about obesity-related molecular changes ^15^. Transcriptomic or proteomic profiles of T2D patients have been studied in tissues including pancreas ^16-20^, adipose tissues ^21,22^, liver ^9,23^, muscle ^11,24,25^, tear samples ^26^, plasma ^27^, and blood vessel organoids ^28^. Similarly, the molecular changes identified across studies varied widely, and only a limited number of genes can be validated in independent cohorts ^19,29^. Therefore, transcriptomic and proteomic profiling of obesity and T2D in humans is far from complete, and further studies are required to systematically evaluate the genetic evidence in support of potential target development programs.

A wide range of animal models have been used to study obesity and T2D, ranging from non-mammalian models such as zebrafish and *Caenorhabditis elegans*, rodent models such as mice and rats, to large animal models such as dogs and pigs as well as nonhuman primates (NHPs) including rhesus macaques and cynomolgus monkeys ^30,31^. Although mice have been the most used preclinical animal model to study metabolic disorders ^31^, they are not as suited for translating findings into clinical applications as NHPs. For example, higher glucose is associated with lower mortality in mice but higher mortality in NHPs and humans, indicating that age-associated metabolic changes in NHPs are more directly translational to humans ^32^.

In contrast to other animal models, NHPs are phylogenetically closest to humans, and their metabolic physiology and anatomy closely resemble those of humans. Therefore, they represent a critical model with unique translational value in biomedical research. Among the commonly used NHP models, the cynomolgus monkey (*Macaca fascicularis*, long-tailed macaque or crab-eating macaque), in particular, has been the most widely used species for drug development ^33^. Although obesity and T2D can be induced by diets in cynomolgus monkeys, they can also develop obesity and T2D spontaneously ^34^, making them an ideal translational model to recapitulate large parts of the pathogenesis of these metabolic diseases in humans. However, the transcriptomic and proteomic profiles of cynomolgus monkey models of obesity and T2D are largely unexplored and therefore poorly understood. To date, only one study has characterized the transcriptome of the liver of type 2 diabetic cynomolgus monkeys induced by high-fat diet ^35^, and no study on the proteome has been reported. Here, we examined the transcriptomic and proteomic profiles of 27 tissues from cynomolgus monkeys to understand the expression patterns in the context of spontaneous obesity and T2D. This extensive and valuable resource of bulk transcriptomes and proteomes across multiple organs enabled us to provide insights into the molecular mechanisms that underlie obesity and T2D.

## RESULTS

### Cynomolgus monkey cohorts and omics data generation

We collected 22 Chinese cynomolgus monkeys including eight obese and four age-matched healthy individuals as well as six type 2 diabetic and four age-matched healthy individuals (Figure 1A, Table S1). All monkeys with obesity or T2D developed the disease spontaneously. BMI was significantly increased in obese monkeys compared with non-obese controls (Figure 1A). T2D monkeys showed significantly increased fasting blood glucose, glycated hemoglobin (HbA1c), triglycerides, and urine glucose when compared with non-T2D monkeys (Figure 1A). Samples from 27 tissues covering all major organs were obtained after necropsy (Figure 1B) and used for profiling the transcriptomes and proteomes, which resulted in 516 transcriptomes and 502 proteomes in the final cohorts after quality control.

**Figure 1.**
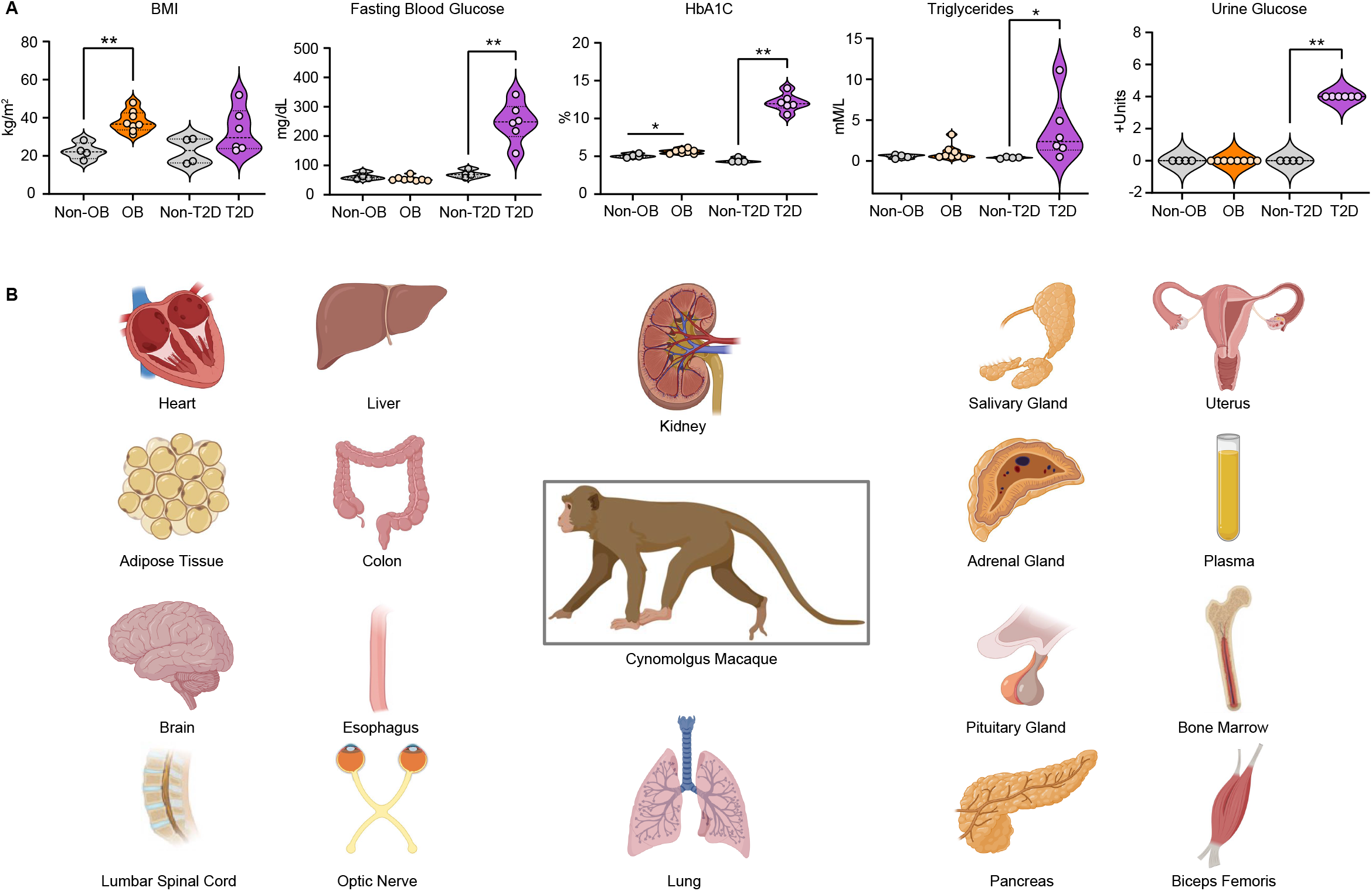
Phenotypes and organs of animals. (A) Violin plots showing body mass index (BMI), fasting blood glucose, glycated hemoglobin (HbA1c), triglycerides, and urine glucose in non-obese (Non-OB), obese (OB), non-diabetic (Non-T2D) and type 2 diabetic (T2D) monkeys. Mann-Whitney U test was performed between diseased and non-diseased groups and statistical significance is denoted as ^*^ p-value < 0.05 and ^**^ < 0.01. (B) Cartoons showing all the major organs collected in the present study. Multiple tissues were included for adipose tissues (abdominal WAT, BAT, visceral WAT), heart (left and right ventricles and atriums), kidney (renal cortex and medulla) and brain (cerebellum, brain stem, thalamus, occipital lobe).

### Transcriptome profiling of obesity across tissues

Hierarchical clustering of the transcriptomes showed tissue-specific expression profiles, and tissues from the same organs exhibited closer correlations than those from different organs (Figure S1A). To interrogate the transcriptional changes associated with obesity, we performed differential expression analysis between obese and non-obese monkeys for all tissues. Adipose tissues showed the largest number of differentially expressed genes (DEGs), while most other tissues showed very few to no DEGs (Figure 2A). Among the three adipose tissues, the largest number of DEGs was found in the abdominal white adipose tissue (WAT, 815 DEGs, Figure S1B), followed by the brown adipose tissue (BAT, 616 DEGs, Figure S1C), while the visceral WAT had the least number of DEGs (206 DEGs, Figure S1D). The DEGs in the BAT and the abdominal WAT were both overrepresented in pathways involving immune response (Figure S1E, Figure 2B). Only one pathway—calcium signaling—was enriched for DEGs in the visceral WAT (false discovery rate [FDR] = 3.02E-04).

**Figure 2.**
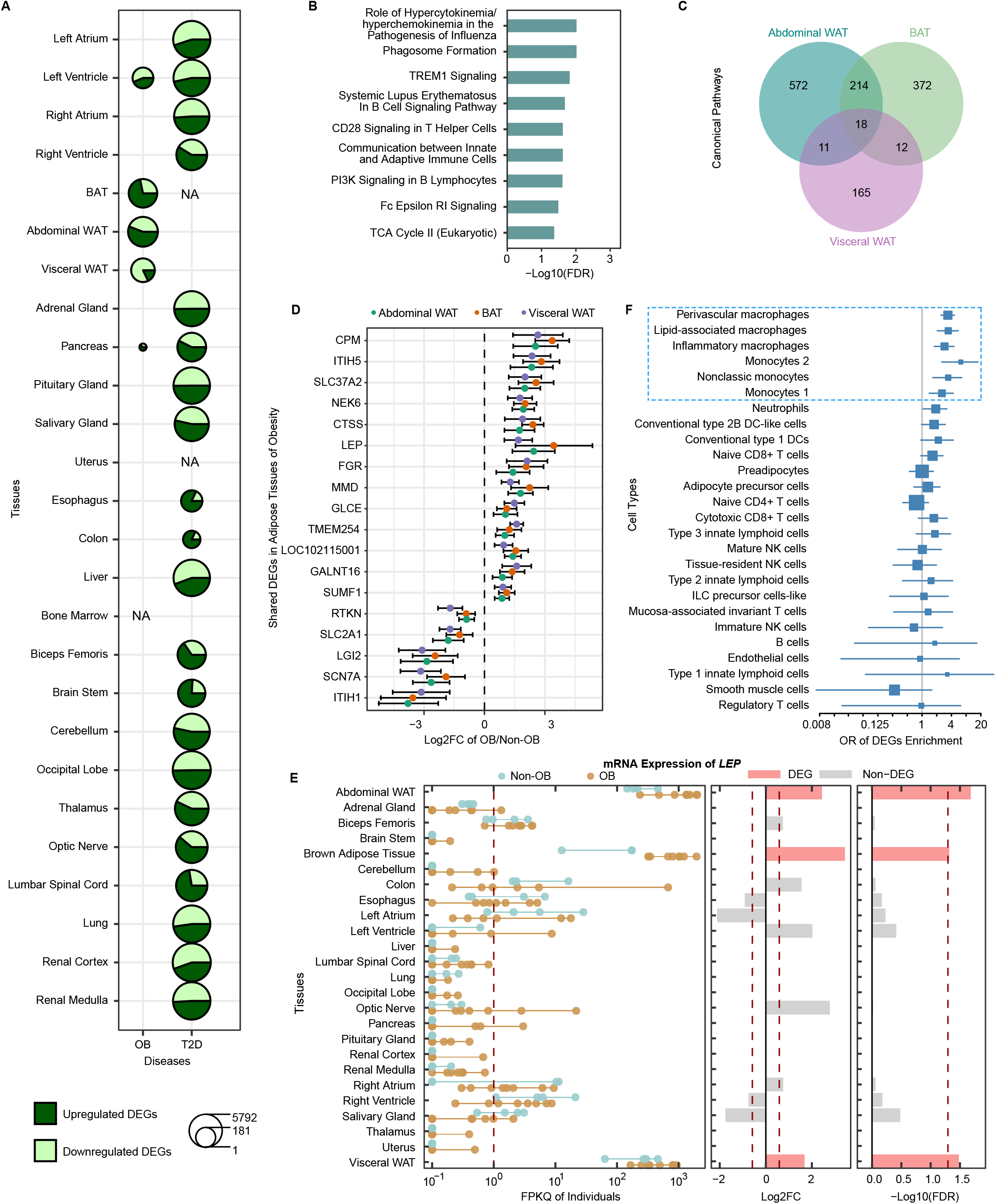
Transcriptomic analyses of obesity. (A) Number of DEGs for obesity and T2D. The size of the circles is proportional to the log2 transformed number of DEGs. (B) Pathways significantly enriched in DEGs of the abdominal WAT in obesity. (C) Venn diagram showing the overlap of DEGs between adipose tissues in obesity. (D) Fold change of the 18 shared DEGs among adipose tissues in obesity. (E) Transcriptional expression of *LEP* across tissues in obesity. The x-axis shows the FPKQ of each sample, log2FC between obese and non-obese monkeys, and -log10 transformed false discovery rates (FDRs; Benjamini-Hochberg corrected p-values) respectively. Differential expression analysis was only performed for tissues with detectable expression (FPKQ > 1). The vertical dashed lines represent the cutoffs for DEGs: FDR < 0.05, log2FC > log2(1.5) or < -log2(1.5). (F) Forest plot showing the enrichment of DEGs of abdominal WAT of obese monkeys in DEGs of various cell types in the stromal vascular fraction of the subcutaneous abdominal WAT from the human obesity single-cell RNA sequencing study. The odds ratio estimates and 95% confidence intervals from Fisher’s exact test for each cell type are shown. The size of the squares is proportional to precision of the estimates. Significant enrichments (FDR < 0.05) are denoted by the blue box. OB, obesity; Non-OB, non-obesity; T2D, type 2 diabetes.

There were 232 genes showing differential expression in both the abdominal WAT and BAT, whereas there were limited number of overlapping DEGs between these two tissues and the visceral WAT (30 and 29, respectively, Figure 2C, Figure S1F). Importantly, all the shared DEGs among the adipose tissues showed same-direction changes. Eighteen genes were differentially expressed between obese vs. non-obese controls in all three adipose tissues including *LEP* and *SLC2A1* (Figure 2C-D). *LEP*, which encodes leptin, plays an important role in energy expenditure and its increased levels have been associated with obesity and T2D ^36^. Our results showed that *LEP* was most abundant in the adipose tissues and was upregulated in all three adipose tissues in obese monkeys (Figure 2E), indicating the shared role of leptin in different types of adipose tissues and its conserved role in obesity between humans and cynomolgus monkeys. *SLC2A1* encodes glucose transporter 1 (GLUT1), which is the major glucose transporter in the mammalian blood-brain barrier ^37^. Previous studies have shown that high-fat feeding suppresses GLUT1 expression in the blood-brain barrier and reduces brain glucose uptake in mice ^38^ as well as induces a decrease of GLUT1 in the subcutaneous adipose tissue in humans ^39^. Higher BMI has been linked to lower *SLC2A1* expression in human subcutaneous adipose tissue ^39^. We observed a significant reduction of *SLC2A1* expression in all three adipose tissues of obese cynomolgus monkeys (Figure S2A), which is consistent with previous studies in mice and humans and thus corroborates the role of

#### *SLC2A1* in obesity

In addition to adipose tissues, 108 DEGs were detected almost uniquely in the left ventricle of the heart (Figure S1F) and not significantly enriched in any canonical pathways. There were five DEGs in the pancreas, including *PSCK1*. Mutations and polymorphisms in *PCSK1* have been strongly associated with human BMI and obesity ^40,41^. We observed a significant increase in *PCSK1*’s expression exclusively in the pancreas of obese monkeys (Figure S2B). Only one DEG each was seen in the cerebellum and the lung, with no DEGs in any other tissues. Collectively, our results suggest that the obesity-related transcriptional changes are mostly limited to adipose tissues.

### Comparison of obesity-associated transcriptional changes between humans and cynomolgus monkeys

To assess whether the transcriptomic profiles of the cynomolgus monkey model of obesity can capture the corresponding signature of human obesity, we compared our dataset with publicly available expression datasets of human obesity (Table S2). For the bulk tissue studies, while there were many DEGs identified in three studies, few to none were seen in others. The number of DEGs varied widely between studies even when the adipose tissue was studied. Therefore, the expression changes in bulk tissues associated with obesity in humans are still inconclusive.

Regardless, we observed 65 shared DEGs showing the same-direction change in the abdominal WAT of obese monkeys with the results from the Lee et al. study ^5^. For the Burkholder et al. study using breast tissue ^12^, we identified 15 overlapping same-direction DEGs in the abdominal WAT. No overlapping DEGs with same-direction changes were detected in the results from other studies. Furthermore, none of these overlapping DEGs were included in the genes associated with obesity in the GWAS catalog. Taken together, expression changes in human obesity need further investigation, and the shared molecular signature between humans and NHPs can be better characterized in the future.

To specifically determine the extent to which obesity-associated expression alterations in human adipose-resident immune cells are observed in cynomolgus monkeys, we evaluated the overlap between the DEGs for each cell type in a recent single-cell RNA sequencing study of obese vs. lean humans using stromal vascular fraction from the subcutaneous abdominal WAT ^42^ and the DEGs identified in the adipose tissues of obese monkeys. For both abdominal WAT and BAT, the largest overlaps were observed for perivascular macrophages (54 and 55 same-direction DEGs, respectively), followed by inflammatory macrophages (24 and 30), which is in line with the frequency alterations of these two macrophage clusters in obese patient samples ^42^. However, no more than two overlapping DEGs were obtained between the visceral WAT of cynomolgus monkeys and any human cell type in that study. Furthermore, the most significant enrichment of DEGs in human macrophages and monocytes was observed for both abdominal WAT (Figure 2F) and BAT, but not visceral WAT of obese monkeys, indicating distinct immune responses in different adipose tissues of obesity. Taken together, our analyses corroborated the role of macrophages and monocytes during human obesity using cynomolgus monkeys, further indicating NHP as a powerful model for studying obesity.

### Proteome profiling of obesity across tissues

The number of detected proteins ranged from 1505 in the biceps femoris to 6140 in the pituitary gland (Figure S3A). Hierarchical clustering of the proteomes showed tissue-specific protein expression profiles (Figure S3B). Differential protein expression analysis between obese and non-obese monkeys identified the largest number of differentially expressed proteins (DEPs) in the adipose tissues: 276 in the abdominal WAT, 151 in the visceral WAT, 186 in the BAT (Figure 3A, Figure S3C-E). Although a smaller number of overlapping DEPs were observed between the adipose tissues compared with DEGs (Figure S3F), the shared DEPs also showed same-direction changes. Four proteins—CD36, ITIH5, LAMB2, and PC—were differentially expressed in all three adipose tissues (Figure S3F). LAMB2 and PC have been associated with waist circumference^43^. CD36 has been linked to high-density lipoprotein (HDL) cholesterol levels ^44^ and plays an important role in lipid absorption ^45^. Pathways overrepresented in DEPs of the abdominal WAT include oxidative phosphorylation, mitochondrial dysfunction, sirtuin signaling pathway, acetyl-CoA biosynthesis I, and fatty acid β-oxidation I (Figure 3B). However, none of these pathways was enriched in DEPs of BAT and visceral WAT. Fewer than 30 DEPs were identified in the cerebellum, uterus, occipital lobe, and pituitary gland, but almost none in other tissues. Taken together, obesity-induced proteomic changes mostly occur in adipose tissues, which is in line with the pattern of transcriptomic data.

**Figure 3.**
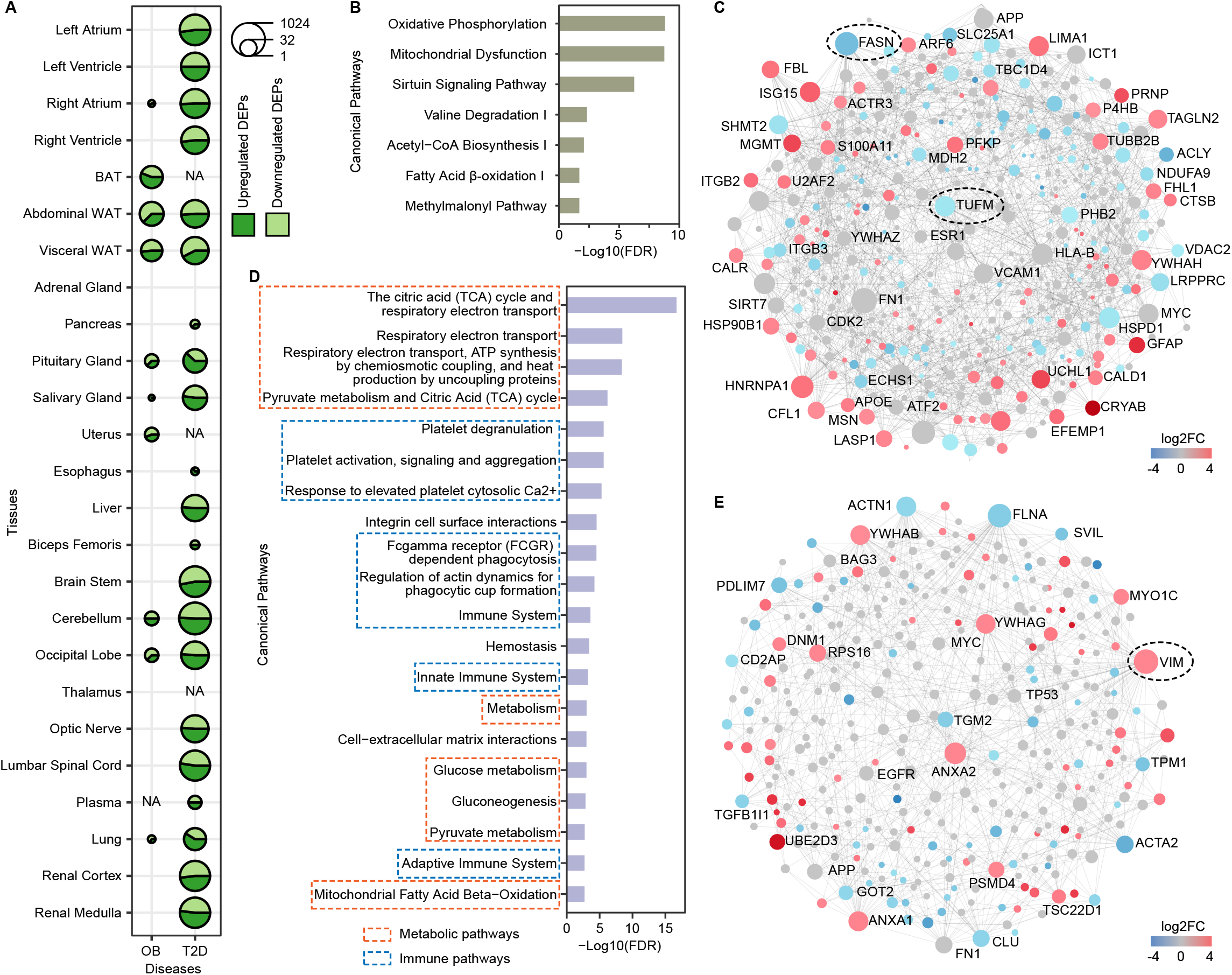
Proteomic analyses of obesity. (A) Number of DEPs for obesity and T2D. The size of the circles is proportional to the log2 transformed number of DEPs. (B) Pathways significantly enriched in DEPs of the abdominal WAT of obesity. (C) Protein-protein interaction network for DEPs of abdominal WAT of obesity. (D) Pathways significantly enriched in the protein-protein interaction network in (C). (E) Protein-protein interaction network for DEPs of visceral WAT of obesity. The size of the circles in the networks is proportional to the number of interactions. Downregulated and upregulated DEPs are in blue and red respectively. OB, obesity; Non-OB, non-obesity; T2D, type 2 diabetes.

### Upstream regulators and secreted factors of adipose tissues in obesity

To determine whether certain upstream regulators drive the proteomic changes associated with obesity in adipose tissues, we sought to identify proteins that were differentially expressed and whose targets were overrepresented in DEPs in obesity. Four proteins—ACLY, ADIPOQ, ITGB3, and THRSP—emerged from the analysis of abdominal WAT. Notably, ACLY and ITGB3 have been associated with waist-to-hip ratio ^43^. ITGB3 has also been implicated in BMI ^46^. THRSP plays a role in the regulation of lipogenesis. ADIPOQ (also known as adiponectin) is secreted by mature adipocytes and can decrease gluconeogenesis in the liver, increase oxidation of fatty acids in the skeletal muscle, and stimulate energy expenditure in the brain ^47^.

Inter-organ crosstalk controlling whole-body metabolism can be coordinated by secreted factors such as ADIPOQ from metabolic tissues, including adipose tissue. Inspired by the finding of ADIPOQ, we sought to identify other factors secreted by adipose tissues in the literature ^47^ that were also differentially expressed in obesity. We found that FABP4 was upregulated in both BAT and visceral WAT, while LPA was downregulated in the visceral WAT. FABP4 is an adipokine that regulates liver lipogenesis and insulin sensitivity in the muscle. Levels of FABP4 have also been reported to be elevated in obese humans ^48^. In contrast, LPA has also been correlated with obesity but appears to negatively impact systemic metabolism ^47^.

### Protein-protein interaction network in the proteome of obesity

To better understand the regulatory networks driving the proteomic changes in obesity, we constructed protein-protein interaction networks using DEPs. For abdominal WAT, 216 of the 276 DEPs were included into one network (Figure 3C). Reactome pathway analysis ^49^ of these genes revealed that metabolic and immune response pathways predominated the network including glucose metabolism (Figure 3D). The most prominent hub protein in terms of the number of interactions was fatty acid synthase—FASN (Figure 3C), which was downregulated in obesity at both mRNA and protein levels. FASN has been associated with BMI ^43^ and proposed as a target for obesity treatment ^50^ and a biomarker of insulin resistance ^51^. Another highly ranked gene, *TUFM*, also showed decreased expression at both mRNA and protein levels and has been linked to BMI ^52^.

The most highly ranked hub gene was *MAPK1* in the BAT and *VIM* in the visceral WAT (Figure S3G, Figure 3E). They have been associated with BMI ^53^ and cholesterol levels ^54^, respectively. The most significant pathway was immune system in both tissues (Figure S3H).

### Comparison of transcriptomic and proteomic alterations in obesity

To examine the relationship between the transcriptome and proteome profiles of obesity, we calculated the correlation of fold changes for DEGs and DEPs. Strong correlation was observed for genes that were both DEGs and DEPs in the abdominal WAT (Figure 4A). There were 82 overlapping DEGs and DEPs in the abdominal WAT (Figure 4B), 27 in the BAT (Figure S4A), and five in the visceral WAT (Figure 4C), but none in any other tissues. All but one of these genes showed same-direction changes at mRNA and protein levels. Only one gene—*ITIH5*—was upregulated in all three adipose tissues. *ITIH5* is a tumor suppressor gene but has also shown increased expression in the human adipose tissue in obesity ^55^. Another gene—*CPM*—was shared between abdominal WAT and visceral WAT, and three other genes—*ATP6V1A, PC*, and *SERPINB9*—were shared between the abdominal WAT and BAT.

**Figure 4.**
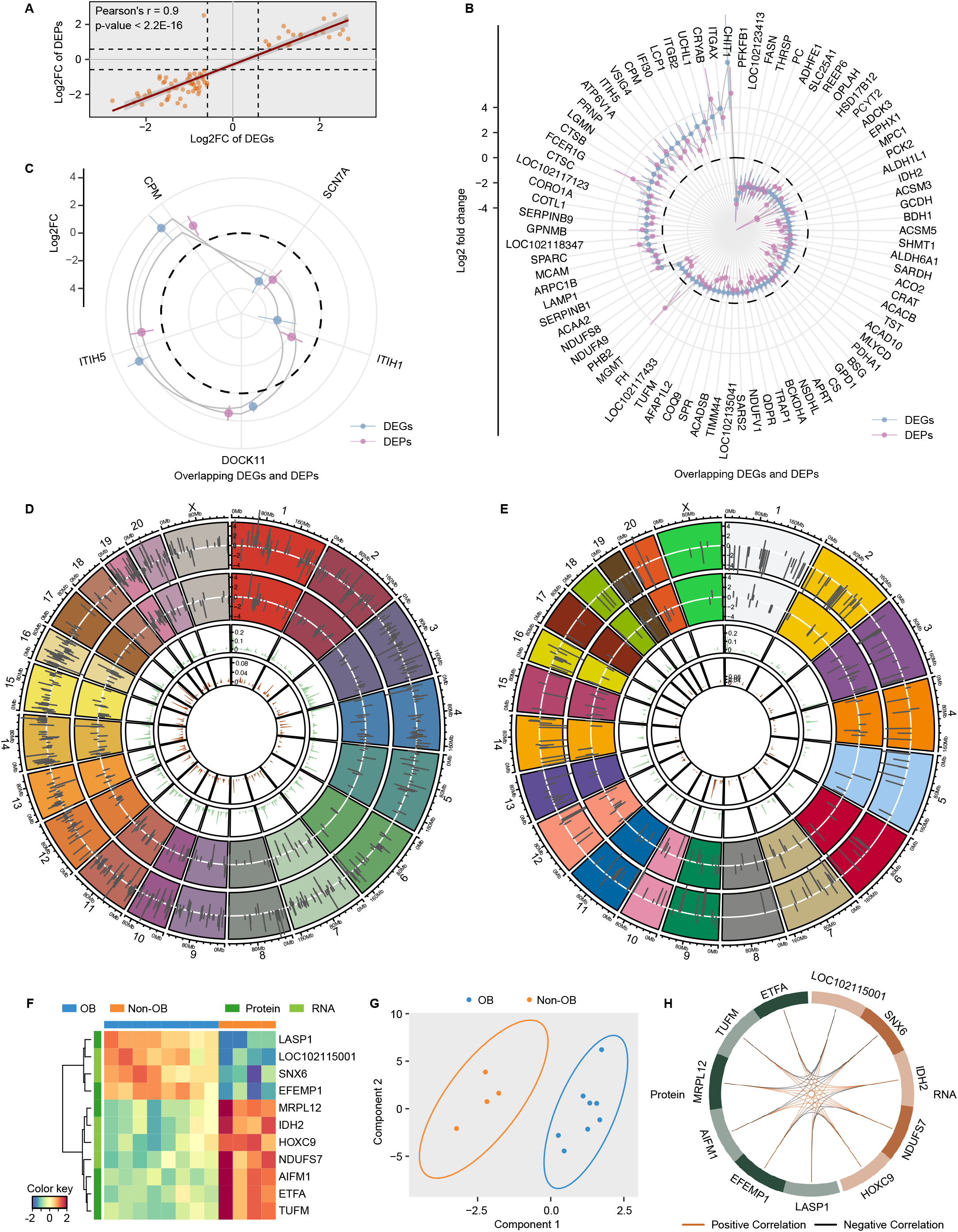
Multi-omics integrative analyses of obesity. (A) Correlation of log2FC of genes that were both DEGs and DEPs in the abdominal WAT of obesity. (B-C) Radar plots showing the 82 and 5 shared DEGs and DEPs in the abdominal and visceral WAT respectively. Error bars represent 95% confidence intervals. (D-E) Circos plots displaying all the DEGs and DEPs genome-wide in the abdominal and visceral WAT, respectively. On each chromosome, log2FCs of DEGs and DEPs and rainfall plots for DEGs and DEPs are vertically ordered from outside to inside tracks. (F) Heatmap of optimal discriminatory molecular signature for the abdominal WAT in obesity identified by the DIABLO method. (G) Consensus plot showing separation of obese and non-obese monkeys using the discriminatory molecular signature for the abdominal WAT. (H) Circos plot showing the positive and negative correlation among the features in the discriminatory molecular signature of the abdominal WAT based on a similarity matrix. Individual tiles on the ring represent individual genes. The connecting lines (orange or black) denote positive or negative correlations between genes with correlation coefficient > 0.7. OB, obesity; Non-OB, non-obesity.

However, most of the DEGs and DEPs in obesity did not overlap in the same adipose tissue (Figure 4D-E, Figure S4B). This indicates that transcriptomic and proteomic perturbations associated with obesity are mostly complementary, illustrating the necessity of studying multi-omics to uncover the complete spectrum of molecular changes of obesity.

### Multi-omics integration for obesity

To shortlist potential multi-omics biomarkers for obesity, we set out to select a minimal and optimal multi-omics signature discriminating obese and non-obese monkeys. We integrated the transcriptomic and proteomic data by applying sparse generalized canonical correlation discriminant analysis via the DIABLO method ^56^. For the abdominal WAT, the multi-omics biomarker panel comprised five mRNAs and six proteins selected from the first component (Figure 4F). These 11 biomarkers were differentially expressed in obesity and can separate the obese and non-obese monkeys (Figure 4G). Strong correlations were observed between these biomarkers (Figure 4H). Notably, six of these 11 genes—*AIFM1, ETFA, IDH2, MRPL12, NDUFS7*, and *TUFM*—play a role in the mitochondria. *IDH2* was downregulated at both mRNA and protein levels in obesity. This gene has been reported to modulate insulin sensitivity and energy expenditure and proposed as a potential therapeutic target for obesity and T2D ^57^. *TUFM* has been associated with BMI ^52^. Other biomarkers in this panel have not been implicated in obesity and may represent novel biomarkers of obesity.

In the BAT, the panel comprised 20 mRNAs and five proteins (Figure S4C), whose expressions were highly correlated (Figure S4D). Moreover, two mRNAs (*CMIP, SNX29*) and two proteins (ATP2A1, HMGB1) are associated with both BMI and T2D in the GWAS catalog. For the visceral WAT, a multi-omics panel of five mRNAs and five proteins was identified (Figure S4E-F). *SMARCC1* is implicated in both BMI and T2D. Altogether, our integrative analysis was able to identify multi-omics biomarker panels that discriminate obese and non-obese monkeys. The panel also included novel molecular features, which provide fresh insights into the molecular ontogeny of obesity.

### Integration of GWAS associations, expression quantitative trait loci (eQTLs), DEGs, and DEPs for obesity

To understand the role of genetic associations for obesity in the transcriptomic and proteomic changes, we obtained a list of 411 genes associated with obesity in the GWAS catalog. No significant enrichment of these genes was seen in the DEGs or DEPs of any tissue. There were only 21 DEGs (Table S3) and eight DEPs (Table S4) associated with obesity, mostly in the adipose tissues. Collectively, our results showed a limited overlap between GWAS findings and our omics study.

Next, we sought to analyze how GWAS signals for obesity can drive gene and protein expression alterations through mediated effects of eQTLs. We first obtained a list of genes (GWAS eGenes) with an overlapped variant of cis-eQTL identified by the GTEx project (www.gtexportal.org) and GWAS loci for obesity in the GWAS catalog for each tissue. Six GWAS eGenes were DEGs (*CTSS, HOXB2, HOXB3, IRS1, PTDSS2, TUFM*) and three were DEPs (TARS2, TUFM, FHL1) in the abdominal WAT, and only one was DEP (SULT1A1) in the visceral WAT. Four of these eGenes—*CTSS, HOXB2, TARS2*, and *TUFM*—were not associated with obesity in the GWAS catalog. Including genes associated with BMI into the analysis identified ten more such genes in the DEPs of the abdominal WAT. For instance, rs12254441 is a variant associated with BMI in the intron of *KAT6B* and thus mapped to *KAT6B* in the GWAS catalog (Figure S5A); however, it is an eQTL of *VDAC2* in the subcutaneous adipose tissue in GTEx, and is in the same topological domain with *VDAC2* in the liver and the pancreas (Figure S5B-C), suggesting that rs12254441 is the link of *VDAC2* with BMI rather than *KAT6B*. Altogether, incorporating eQTLs into the analysis has the potential to uncover additional obesity-associated genes.

A recent large-scale human exome sequencing study identified 16 genes significantly associated with BMI for the burden of rare nonsynonymous variants ^58^. Five genes were DEGs in our study, which results in significant enrichment (Fisher’s exact test p-value = 8.01E-03, odds ratio = 4.83). *UHMK1* was upregulated, and *CALCR* and *PDE3B* were downregulated in the abdominal WAT. *PCSK1* was upregulated in the pancreas. *SPARC* was upregulated in both abdominal WAT and BAT. Additionally, SPARC was an upregulated DEP in both abdominal and visceral WAT. We envision that these five genes with both human genetic evidence and multi-omics validation can be prioritized for effective therapeutic strategies.

### Transcriptome and proteome profiling of T2D across tissues

To dissect the molecular mechanism of T2D, we carried out differential expression analysis between T2D vs. non-diabetic controls at both transcriptomic and proteomic levels. Unlike in obesity, we observed extensive transcriptomic and proteomic perturbations across most tissues (Figure 2A, Figure 3A). T2D is mainly characterized by impaired insulin secretion by pancreatic β cells. In the pancreas, 646 DEGs were identified, and the most significant pathway enriched was Gαq signaling (FDR = 4.35E-02).

In contrast, we found only eight DEPs—insulin, ECHDC2, ARL6IP1, GON7, CISD3, SFRP1, SIAE, and COL6A1—in the pancreas. As expected, insulin was the most significantly decreased protein in T2D (FDR = 3.51E-02, log2 fold change [log2FC] = -4.33), indicating that cynomolgus monkeys are a valid model for studying T2D. Another DEP—CISD3—was also differentially expressed at the transcriptional level. However, no associations between these DEPs and T2D have been reported previously, except for insulin, illustrating that novel associations can be detected using cynomolgus monkeys as a model. There were 22 DEPs in the plasma, including three proteins (LPA, CETP, and APOC4) involved in lipid metabolism. We observed decreased expression of LPA and CETP but increased expression of APOC4 in T2D. Previous studies have reported associations between low levels of LPA and increased risk of T2D ^59^.

We next focused on the genes that were both DEGs and DEPs and showed same-direction changes at mRNA and protein levels (same-direction DEG/DEPs) in T2D compared with non-diabetic controls. Individuals with T2D are susceptible to complications such as kidney and cardiovascular diseases. Intriguingly, tissues from the kidney and heart were also among tissues with the highest number of these genes. Two kidney tissues—renal medulla and cortex—had the highest number of same-direction DEG/DEPs (288 and 255, respectively) among all tissues, followed by two brain tissues (cerebellum and occipital lobe) and two left heart tissues, and then the liver (Figure 5A). Insulin resistance in T2D can be present not only in pancreatic β cells but also in tissues including liver, muscle, and brain ^60^, which is consistent with the relatively high number of same-direction DEG/DEPs in liver and brain tissues such as the occipital lobe and cerebellum (Figure 5A), although there was no such gene in the biceps femoris. There were a medium number of same-direction DEG/DEPs in nerve tissues. Additionally, two endocrine tissues—pituitary gland and adrenal gland—had 50 and 46 same-direction DEG/DEPs, respectively. Few to no DEGs were seen in other tissues.

**Figure 5.**
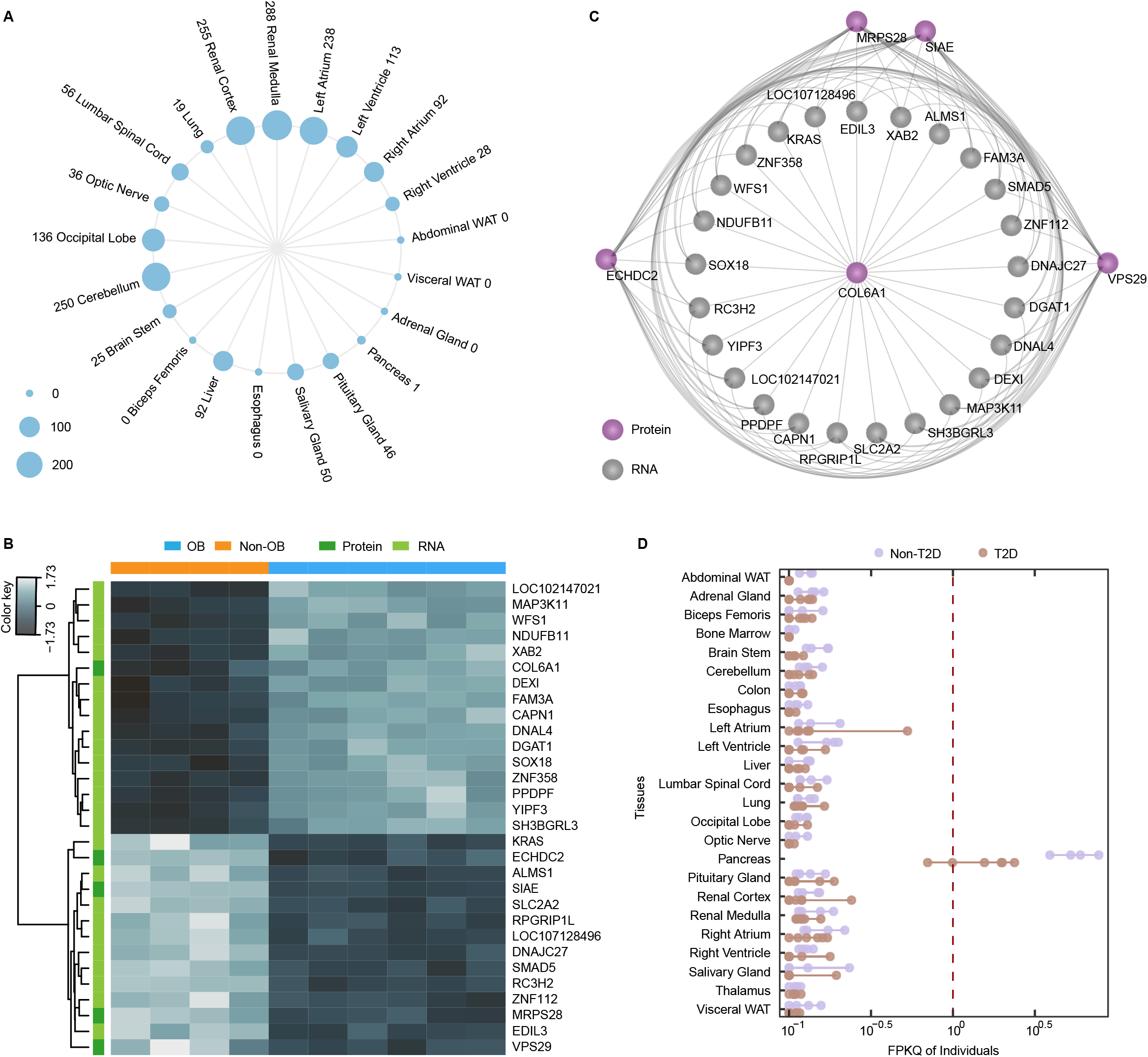
Multi-omics analyses of T2D. (A) Circular dot plot showing the number of shared same-direction DEGs/DEPs across tissues for T2D. (B) Heatmap of optimal discriminatory molecular signature for the pancreas in T2D identified by the DIABLO method. (C) Relevance network displaying the correlation among the features (proteins in pink circles and RNAs in grey circles) in (B). Features with a correlation coefficient > 0.7 are connected by lines. (D) Transcriptional expression of *SLC30A8* across tissues in T2D. Each point represents a sample and colors denote the groups. OB, obesity; Non-OB, non-obesity; T2D, type 2 diabetes; Non-T2D, non-diabetes.

### Multi-omics integration for T2D

We also applied the DIABLO method to T2D to look for a multi-omics biomarker panel distinguishing the diseased and healthy monkeys. For the pancreas, a panel comprising 25 mRNAs and 5 proteins was selected (Figure 5B). Protein biomarkers were highly correlated with the mRNA biomarkers (Figure 5C). *SLC2A2* is associated with both T2D and obesity, and *DNAJC27* is associated with both obesity and BMI in the GWAS catalog. *MAP3K11, RPGRIP1L*, and *WFS1* are implicated in T2D, and *PPDPF, RPGRIP1L*, and *RC3H2* are linked to BMI. Our analysis selected discriminant and well correlated molecular features across omics data types, which provides potential biomarkers of T2D for validation in future studies.

### Integration of GWAS associations, eQTLs, DEGs, and DEPs for T2D

To identify the genes with GWAS associations in DEGs and DEPs, we curated a list of 2009 genes associated with T2D from the GWAS catalog. There were 639 and 388 GWAS genes overlapping with DEGs and DEPs, respectively, of which 194 and 155 were DEGs or DEPs in only one tissue. GWAS genes seemed more likely to be differentially expressed at the transcriptional level in at least one tissue compared with non-GWAS genes (Fisher’s exact test p-value = 8.13E-04, odds ratio = 1.23) but not at the proteomic level. There were 49 DEGs with GWAS signals in the pancreas, including *SLC30A8* and *SLC2A2*. These two genes have been associated with T2D in multiple studies ^61,62^. Importantly, *SLC30A8*, that was exclusively expressed in the pancreas, was significantly downregulated in T2D monkeys (FDR = 4.91E-03, log2FC = -2.05; Figure 5D) due to the disease-related damage and necrosis of Langerhans Islets. *SLC2A2* had detectable expression in only several tissues (Figure S5D) and showed significantly lower expression in the pancreas (ranked second among all genes, FDR = 1.16E-03, log2FC = -3.76), liver (FDR = 6.48E-04, log2FC = -1.16), and renal medulla (FDR = 1.39E-04, log2FC = -5.06) of T2D individuals. Note that the downregulation of *SLC2A2* in human patients with T2D has been replicated in three different cohorts by previous work ^19^. Moreover, insulin was the only DEP in pancreas with GWAS evidence. Altogether, our results support the association of GWAS genes such as *SLC30A8, SLC2A2*, and insulin with T2D and suggest that cynomolgus monkeys can be a powerful animal model to study the effects of potential treatments targeting these genes.

Among the same-direction DEGs/DEPs, we identified at least one T2D GWAS gene in 14 tissues, including 19 in the renal cortex, 16 in the left atrium, and 12 in the cerebellum (Table S5). The finding of these genes in multiple tissues demonstrates that T2D is not a disease damaging the pancreas only, but rather a group of conditions and complications throughout the body. These results can be used to prioritize biologically meaningful biomarkers and therapeutic targets for T2D and its complications. For example, *GNPDA2* was downregulated at both mRNA and protein levels in both the kidney and heart and has also been associated with obesity. Three genes—*LPL, APOE*, and *G6PD*—are involved in lipid metabolism and exhibited decreased expression in the ventricles of the heart and increased expression in the liver and the cerebellum, respectively. Importantly, two genes—*PIK3R1* and *RPTOR*—involved in insulin-related pathways were differentially expressed in the pituitary gland, indicating that other endocrine organs in addition to the pancreas can also play an important role in T2D.

Next, we evaluated the overlap of GWAS eGenes of T2D and DEGs or DEPs in T2D in the same manner as we analyzed the dataset of obesity. We found differentially expressed GWAS eGenes at the transcriptional level in 12 tissues, including nine in the pancreas, among which four genes— *RPL13, SPPL2A, TH*, and *UBLCP1*—were included in the GWAS catalog, and the other five genes were not (*PAK4, SH2B1, MAPK3, MAN2C1, YPEL3*). Our results demonstrate that incorporating e-QTLs into the analysis can identify additional potential causal genes of T2D. For example, rs8046545 is a T2D-asssociated variant in the intron of *ATP2A1*; however, this variant is a cis-eQTL of the gene *SH2B1* in the pancreas, indicating that this GWAS signal might point to *SH2B1* rather than *ATP2A1*. Indeed, decreased levels of *SH2B1* result in both leptin and insulin resistance and thus obesity and diabetes in mice ^63^. Previous functional studies have also reported that *SH2B1* can regulate glucose metabolism and promote insulin/insulin-like signaling in mammals ^64,65^. At the proteomic level, we found differentially expressed GWAS eGenes in nine tissues, including 20 in abdominal WAT but none in the pancreas.

### Comparison of T2D-associated transcriptional changes between humans and cynomolgus monkeys

Previous studies have described the expression profile of T2D in several human tissues. However, even when the same tissue was used, the number of DEGs identified varied across datasets and the overlap was generally poor between studies. For example, a previous study leveraging the transcriptomes of three cohorts of type 2 diabetics pinpointed only three genes shared across cohorts ^19^; another study reanalyzing transcriptomes of adipose tissue from two independent cohorts of patients with T2D found only 15 shared downregulated DEGs ^29^.

Regardless, we compared our results with three human pancreatic islet transcriptomic datasets of T2D: GSE41762 ^17^, GSE38642 ^18^, and GSE50244 ^16^. There were only 13 DEGs in GSE41762 (at the threshold of FDR < 0.05 and |log2FC| > log2(1.5)) and six DEGs in GSE38462 (FDR < 0.1 and |log2FC| > log2(1.5)). The DEGs of pancreas in our study shared only one DEG—*SLC2A2*— with GSE41762, but no overlap was found with GSE38642. There were 1390 DEGs in GSE50244 (FDR < 0.05 and |log2FC| > log2(1.5)), of which 102 DEGs overlapped with the DEGs of pancreas in cynomolgus monkeys. Almost all these shared DEGs (100 of 102) showed same-direction change in both studies, and ten of them were with GWAS signals, including *SLC2A2* (Table S6). Overall, whereas characterization of transcriptome changes linked to T2D in humans needs further investigation, we were able to prioritize a subset of genes such as *SLC2A2* with multiple lines of evidence as potential T2D drug targets.

### Shared transcriptional changes between obesity and T2D

Considering obesity is one of the critical risk factors of T2D, we attempted to determine to what extent cynomolgus monkeys with obesity and T2D shared similar transcriptional changes. There were two shared DEGs—*GAD2* and *LOC102137725*—in the pancreas between obesity and T2D. *GAD2* was upregulated in obesity but downregulated in T2D. It is a major autoantigen in insulin-dependent diabetes and has been associated with lean body mass ^66^.

We next analyzed the expression profiles of the two adipose tissues—abdominal and visceral WATs—available for both groups using weighted-gene co-expression network analysis (WGCNA) ^67^ to infer functionally related genes in the form of co-expression modules associated with both diseases. We identified 23 modules in the abdominal WAT, of which four were significantly associated with both obesity and T2D (Figure S6A) after multiple testing correction. The first module (green module) consisted of 1075 upregulated genes in obesity and T2D (Figure 6A). These genes were overrepresented in many pathways involved in immune response (Figure 6B). All the other three modules contained downregulated genes in obesity and T2D. Interestingly, the brown module consisting of 1653 genes was enriched in metabolism-related pathways such as fatty acid β-oxidation, with oxidative phosphorylation and mitochondrial dysfunction being the most significant ones (Figure 6C).

**Figure 6.**
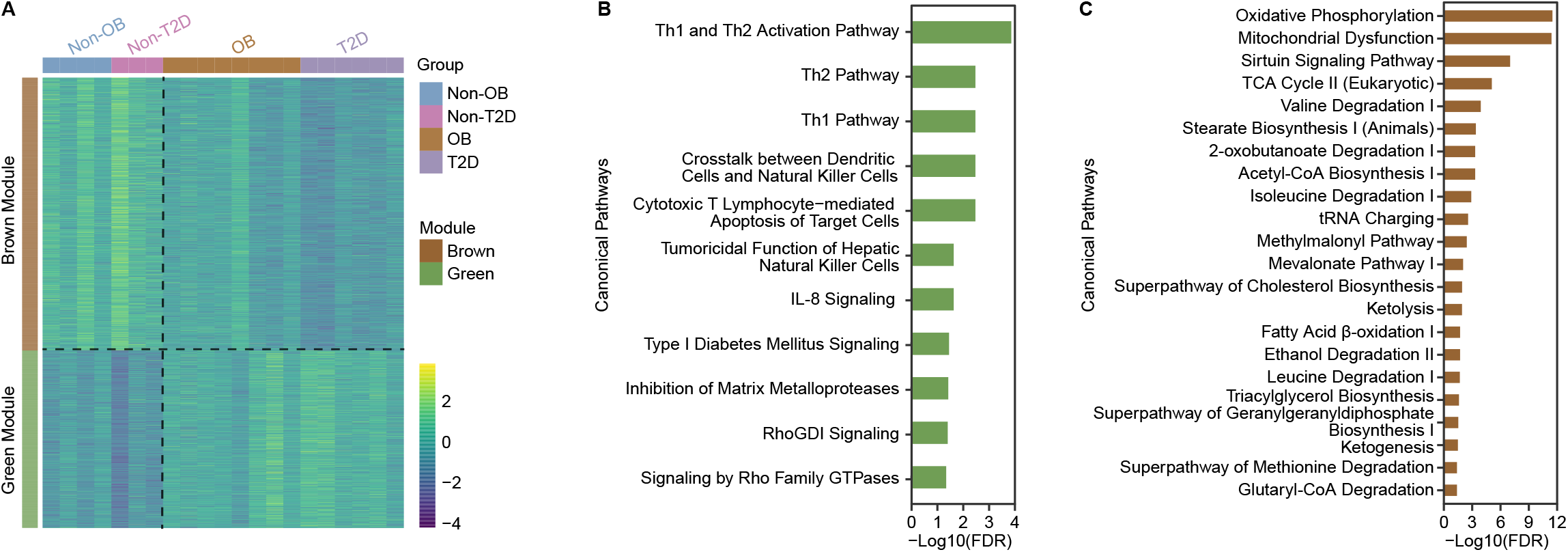
Shared transcriptional alterations in the abdominal WAT between obesity and T2D. (A) Heatmap showing expression of the genes in the green and brown modules identified by WGCNA. Color bars indicate different modules and groups. Genes in the brown module were downregulated in obesity and T2D but genes in the green module were upregulated. (B) Significant pathways overrepresented in the genes of the green module. (C) Significant pathways overrepresented in the genes of the brown module. OB, obesity; Non-OB, non-obesity; T2D, type 2 diabetes; Non-T2D, non-diabetes; WGCNA, weighted-gene co-expression network analysis.

Of the 26 co-expression modules inferred from the visceral WAT, two were associated with both obesity and T2D (Figure S6B-C). The first module (purple module) contained 722 upregulated genes in the two diseases (Figure S6C) and was enriched in many signaling pathways, including leukocyte extravasation signaling pathway (Figure S6D), which is involved in the process by which leukocytes migrate from the blood to the tissue during inflammation. The second module (darkred module) had 130 downregulated genes (Figure S6C) including three genes—*EML6, HS6ST3*, and *LINGO2*—with GWAS signals for both obesity and T2D.

### Comparison of proteomic changes between obesity and T2D

To address the proteomic features associated with the disease progression from obesity to T2D, we first compared the DEPs within the same tissues between these two diseases. Whereas most DEPs appeared to be unique to one disease, we observed 71 shared DEPs in the abdominal WAT (Figure 7A), 21 in the visceral WAT (Figure 7B), 10 in the cerebellum, and few to none in other tissues (Table S7). In the abdominal WAT, 62 of the 71 shared DEPs showed same-direction changes between obesity and T2D (Figure 7C). These genes were enriched in metabolic pathways such as acetyl-CoA biosynthesis (Figure S6E). Four of these genes—*PCCB, CHDH, SERPINB1*, and *ITGAX*—are associated with BMI, and another three genes—*LAMB2, PPIF*, and *GCDH*—are linked to T2D. In the visceral WAT, there were only seven shared DEPs with same-direction changes between obesity and T2D, suggesting that the visceral WAT may undergo more extensive proteomic dysregulations when progressing from obesity to T2D than the abdominal WAT. Furthermore, *ITLN1* was downregulated in obesity but upregulated in T2D in the pituitary gland. *ITLN1* may function as an adipokine enhancing insulin-stimulated glucose uptake in adipocytes, which is also in line with our observation that it was upregulated in the visceral WAT of T2D.

**Figure 7.**
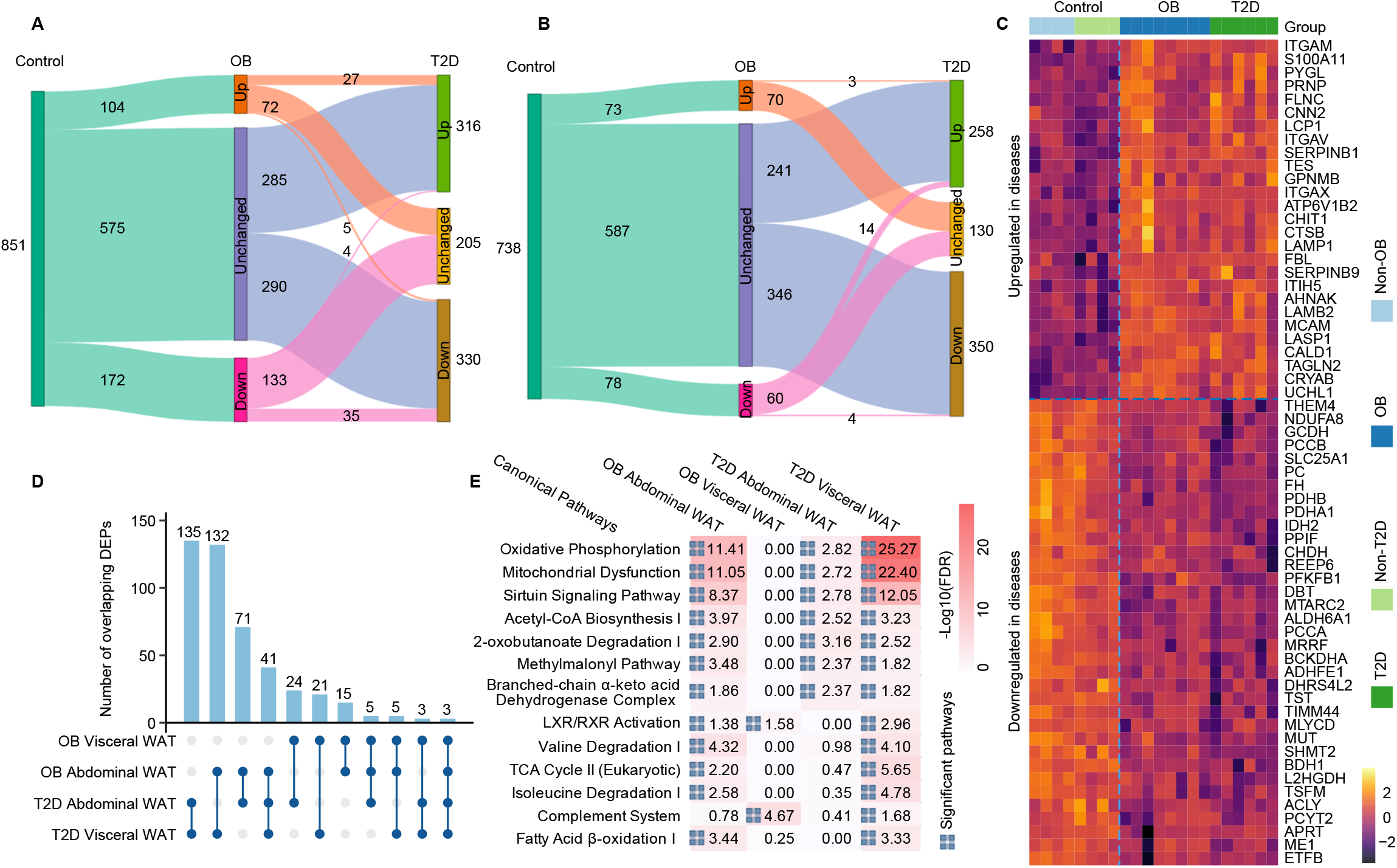
Comparison of proteomic alterations in the adipose tissues between obesity and T2D. (A-B) Sankey diagrams showing the change of DEPs in the abdominal and visceral WAT between obesity and T2D, respectively. Upregulated and downregulated DEPs are labeled as “Up” and “Down,” respectively. Proteins without changes are labeled as “Unchanged”. Only the DEPs in at least one disease were counted. (C) Expression of the 62 DEPs showing same-direction changes in the abdominal WAT between obesity and T2D. The horizontal dashed line separates the upregulated and downregulated DEPs in diseases whereas the vertical one separates the control and disease groups. (D) UpSet plot showing the number of overlapping DEPs between the abdominal and visceral WATs of obesity and T2D. (E) Pathways enriched in DEPs of the abdominal and the visceral WAT in obesity and T2D. The values in the cells are -log10 transformed FDRs. The squares denote significant pathways.

We next compared the DEPs across adipose tissues and diseases. Whereas the visceral WAT of obesity showed a limited overlap of DEPs (24 proteins) with the abdominal WAT of T2D, the visceral WAT of T2D shared a much larger number of DEPs (132 proteins) with the abdominal WAT of obesity, of which all but one exhibited same-direction changes (Figure 7D). Intriguingly, the top pathways enriched in DEPs in the visceral WAT of T2D were almost the same as in the abdominal WAT of obesity but only had two overlaps with the visceral WAT of obesity (Figure 7E), indicating that the visceral WAT might shift part of its metabolic activities toward the abdominal WAT when progressing from obesity to T2D. Three genes—*GPNMB, PC*, and *PCYT2*—showed same-direction differential protein expression in both adipose tissues and both diseases. None of them has been associated with obesity or T2D. Notably, *PC* encodes pyruvate carboxylase, which is involved in gluconeogenesis, lipogenesis, and insulin secretion.

## DISCUSSION

We have quantitatively analyzed the transcriptomes and proteomes of cynomolgus monkeys with spontaneous obesity and T2D together with age-matched healthy controls. While characterization of genetic effects is still limited by sample size, our study represents a deep survey of molecular profiles of obesity and T2D across many tissues in NHP models.

Using multi-tissue transcriptomes and proteomes, we were able to establish that the gene expression alterations linked to obesity were mainly limited to adipose tissues. Most obesity-associated genes identified by GWAS were not differentially expressed in obese monkeys. Future larger GWAS and multi-omics studies are likely to uncover many more obesity-associated genes. It is possible that post-transcriptional and post-translational modifications as well as inter-species differences may also contribute to the underrepresentation of obesity GWAS genes in the DEGs and DEPs detected in our study. Despite these limitations, our integrative analysis still identified certain putatively causal genes (Table S3, Table S4). Note that the discrepancy between the transcriptional changes and GWAS findings has been previously reported for both obesity and T2D ^68^. It has also been proposed that gene expression changes and genetic associations from GWAS may cover different biological aspects of the diseases ^68^, which underscores the importance of the present study to discover more potential therapeutic targets in addition to GWAS. For example, *ISM1* was downregulated in both obese and type 2 diabetic cynomolgus monkeys (Figure S7A-B) but is not implicated in obesity and T2D by GWAS. However, *ISM1* has been recently identified as an adipokine playing a dual role in increasing adipose glucose uptake while suppressing hepatic lipid synthesis and has been shown to have therapeutic potential for diabetes and hepatic steatosis ^69^. Future research will need to examine the effects of such possible targets without GWAS support for pharmacological interventions in metabolic diseases.

Our study provides an open resource to prioritize potential new targets and pathways for obesity. For example, we envision that further investigation into the DEGs might yield potential drug discoveries in humans, especially for the five DEGs with rare coding variant associations ^58^ such as *SPARC* and *PDE3B*, as well as the 18 DEGs and four DEPs shared among all adipose tissues such as *SLC2A1* and *CD36*. DEPs in other tissues such as PGM2L1 (glucose 1,6-bisphosphate synthase) in the occipital lobe might be worth further investigation as well. Sirtuin signaling pathway was upregulated in the abdominal WAT in obese monkeys at the protein level but it has not been well studied in obesity. Substantiating the role and elucidating the mechanism of this pathway in obesity may also inform future targeted therapies.

In contrast to obesity, we observed molecular changes in most tissues in T2D. Although insulin secretion by the pancreas is one of the core defects in T2D, there was a relatively small number of DEGs and DEPs in pancreas compared with many other tissues including kidney, heart, and brain. The monkeys with T2D in our cohorts showed symptoms of kidney disease, although we did not have ample clinical evidence to distinguish between diabetes complications and non-diabetic kidney diseases. Regardless, the large-scale expression perturbations in kidney tissues may arise partly owing to the kidney damage in monkeys with T2D. However, the diabetic monkeys in our study did not manifest complications in the heart and brain, indicating that the molecular changes in these organs could have occurred prior to or independently of the diabetic complications. Furthermore, the expression perturbations in many tissues may also exist partly owing to the inter-organ communication via secreted factors such as leptin and adiponectin in coordinating whole-body metabolism, through which one tissue can affect metabolism in physically distant tissues. Previous studies on human T2D have mostly focused on single tissues, which lacked the power of characterizing a comprehensive map of T2D-related molecular profiles across organs. Moreover, we highlighted some potential drug targets for T2D in our study, such as *SLC30A8* and *SLC2A2*, as well as pathways such as Gαq signaling. Gαq is a member of α-subunits of G proteins, which can initiate various intracellular signaling cascades. Gq-coupled receptors have been reported to play an important role in regulating hepatic glucose fluxes and suggested to be novel receptor targets for the treatment of T2D ^70^. Moreover, activation of Gq signaling can stimulate glucose uptake in human skeletal muscle cells ^71^. Together, our study leveraging multi-tissue multi-omics revealed that T2D could develop into a multifactorial, complex, and holistic disease impacting various organs as the disease progresses and provides a valuable resource for understanding the molecular underpinnings of T2D.

Our co-expression analysis of transcriptomics data captured shared modules of genes associated with both obesity and T2D in the adipose tissues, supporting that obesity can contribute to the development of T2D at the transcription level. Importantly, immune response-related pathways were upregulated whereas the lipid metabolism and mitochondrial pathways were downregulated, which is in line with previous findings in human and rodent models ^8,60,72^. Indeed, the adipose tissues in obesity and diabetes are infiltrated with immune cells and increase secretion of pro-inflammatory cytokines ^47^. Previous studies also provided the evidence that dysregulation of lipid metabolism and altered mitochondrial function are closely associated with T2D ^47^. At the proteomic level, we also observed shared DEPs between the two diseases (Figure 7A-B). Interfering with these shared molecular alterations, such as *GAD2* in the pancreas and *LAMB2* in the adipose tissues, might help with blocking or slowing the disease progression from obesity to T2D.

On the other hand, we discovered some genes showing divergent changes in expression between obesity and T2D. For example, one of the widely reported genes implicated in T2D—*SLC30A8*— was upregulated in the pancreas in obese monkeys but downregulated in diabetic monkeys at the transcriptional level (Figure S7C, Figure 5D). CTSC and several mitochondrial proteins (ATP5MG, IBA57, and TOMM6) also showed an opposite direction of alterations. Furthermore, most of the DEPs were only observed in one disease. The visceral WAT exhibited distinct molecular signatures in obesity and T2D and underwent major changes when progressing from obesity to T2D. Our findings demonstrate the importance of targeting these genes at various stages of disease progression.

Our study illustrates that deciphering the exhaustive molecular underpinnings of obesity and T2D necessitates complementary multi-omics studies. Most perturbed genes only showed differential expression in diseased monkeys at either the transcriptional or protein level, and the impacted pathways were also unique to one type of omics data, suggesting that molecular alterations associated with obesity and T2D at various molecular levels were mostly complementary rather than convergent.

To conclude, we profiled the bulk transcriptomes and proteomes of all major organs across multiple cynomolgus monkeys to create a comprehensive resource and demonstrated that NHPs are a powerful model for studying obesity and T2D. Although all currently available animal models of obesity and T2D have distinct advantages and limitations, NHP models, especially cynomolgus monkeys, remain indispensable for therapeutic discovery and optimization to treat these diseases in humans due to their close genetic relationship with and physiological similarity to humans. Future endeavors such as single-cell RNA sequencing will further advance our understanding of the molecular signatures of obesity and T2D at a higher resolution.

## Acknowledgments

We thank Dr. Christine Karbowski at Amgen for valuable discussions and thank Shanghai OE Biotechnology Company and Shanghai Biotechnology Corporation for their valuable contributions to DIA-MS and RNA-seq processing, respectively.

## Author Contributions

Y.-H.H., L.G.H., M.M.V., and S.W. designed the study. X.Z. performed the analyses and wrote the manuscript. Y.L. conducted the experiments. M.S. analyzed the phenotypic data and contributed to the study design. O.H. contributed to data processing.

## Methods

### Data availability

All RNA-seq data generated in this study have been deposited in the GEO under accession number GSE188418.

### Ethical statement

All the animals used in this study and the experimental protocols and procedures were approved by the Institutional Animal Care and Use Committee of WuXi AppTec (Suzhou, China) Co., Ltd. and JOINN Laboratories (Suzhou, China) Co., Ltd., in accordance with the guidelines of Association for Assessment and Accreditation of Laboratory Animal Care.

### Animal collection

Spontaneous obesity in cynomolgus monkeys were defined as BMI ≥ 30 kg/m^2^. For T2D, the American Diabetes Association criteria for diabetes are fasting blood glucose (FBG) ≥ 126 mg/dL and HbA1c ≥ 6.5% in humans. However, the FBG is typically ∼20 mg/dL lower in normal NHPs than in healthy humans, and the mean HbA1c in NHPs is ≤ 5.0%. All cynomolgus monkeys with spontaneous T2D collected in this study had FBG ≥ 140 mg/dL and HbA1c > 10%. Age-matched healthy monkeys were also collected. Twenty-seven tissues for each animal were dissected and snap-frozen after necropsy.

### RNA extraction, cDNA library preparation, and Illumina sequencing

Total RNA was extracted from 27 snap-froze tissues with RNeasy Micro/Mini Kit (QIAGEN, Germany, Cat. #74004/#74106), according to the manufacturer’s instructions. The quality of the isolated RNA was analyzed by Agilent Bioanalyzer 2100. Only the RNA samples with an RNA Integrity Number (RIN) value of 6.0 or higher were included for RNA sequencing. RNA-seq libraries were constructed from RNA samples meeting the QC criteria based on polyA+ selection of mRNA using a standard strand-specific protocol with VAHTS^®^ Stranded mRNA-seq Library Prep Kit for Illumina (Vazyme, China, Cat. #NR612). The quality of the sequencing libraries was analyzed by Agilent Bioanalyzer 4200 and the peak length ranged from 250 to 500 bp. Clusters were generated by cBot with the library diluted to 10 pM and then sequenced on Illumina HiSeq 2500 instruments with 2 × 150 bp paired-end sequencing. All libraries were sequenced to an average depth of 52 ± 8.8 (mean ± SD) million reads (corresponding to 26 ± 4.4 million paired-end reads).

### RNA-seq processing

The initial analysis of RNA-seq data was processed by our internal pipeline involving read filtering and trimming, alignment, quantification, and normalization. All read files were processed by fastp (v0.19.8) ^73^ to remove poor quality reads and to trim poor quality 3’ bases. The filtered and trimmed reads were aligned and quantified using OmicSoft’s (Cary, NC, USA) Array Studio software (https://www.omicsoft.com/publish-studio; Oshell.exe v10.0) to reference genome NCBI Macaca_fascicularis_5.0. The transcriptomic annotation NCBI Macaca fascicularis Annotation Release 101 was used as the transcript model reference. The alignment algorithm OSA ^74^ is documented in an OmicSoft whitepaper (http://omicsoft.com/downloads/whitepaper/OmicsoftAligner.pdf), and the gene/transcript quantification module is an OmicSoft implementation of the RSEM algorithm ^75^.

Single nucleotide polymorphisms were called from the RNA-seq data and the Pearson correlation was calculated between each sample’s mutation frequencies. Samples with mis-assigned donors were excluded from further analysis. Pearson correlation was also calculated between each sample’s expression profiles and samples with mis-assigned tissues were removed from the dataset.

Only genes with fragments per kilobase million quantile normalized (FPKQ) ≥ 1 in at least one of the two groups in the same comparison were included for further statistical testing. Differential expression analysis was performed using limma voom (v3.42.2) ^76^ after quantile normalization. Genes with Benjamini-Hochberg corrected p-value < 0.05 (for the comparison of obese and non-obese controls) or < 0.01 (for the comparison of T2D and non-diabetic controls) and more than 1.5-fold change (|log2FC| > log2(1.5)) were chosen as DEGs.

### Co-expression analysis

WGCNA (v 1.46) ^67^ was used to identify modules of co-expressed genes using all the genes with detectable expression (total read counts > 100). Quantile normalized count data obtained from the “voom” function in limma ^76^ was used for analysis. Signed network was constructed.

### Protein extraction and digestion

The same 27 tissue types for RNA isolation were used for protein extraction. Snap-frozen tissues were lysed in SDS lysis buffer (Beyotime, China, Cat. #P0013G) supplemented with 1 mM PMSF (Amresco, USA, Cat. #PB0425). Cells were disrupted by sonication on ice for 3 min, followed by centrifugation at 12,000 g for 10 min at 4°C. Then concentration of the protein was measured by the Pierce™ BCA Protein Assay Kit (Thermo Fisher Scientific, USA, Cat. #23227). Samples were stored at -80°C until proteomic preparation.

For each sample, 30 μg protein was reduced at 60 °C with reducing buffer (8 M urea, 100 mM TEAB, 10 mM DTT, pH 8.0), alkylated with iodoacetamide (Sangong Biotech, China, Cat. #A600539), and digested with trypsin (Hualishi Scientific, China, Cat. #HLS TRY001C) overnight, followed by desalting with SOLA™ SPE plates (Thermo Fisher Scientific, USA, Cat. #60309-001). Samples were vacuum-evaporated and resuspended in 0.1% formic acid in ddH_2_O and iRT peptides (Biognosys, Switzerland, Cat. #Ki-3002) were added (1:10).

### Data-dependent and data-independent acquisition (DDA and DIA) mass spectrometry (MS)

Peptides were separated on an 1100 HPLC System (Agilent, USA) connected to an Agilent ZORBAX Extend-C18 column (80 Å, 150 mm × 2.1 mm, 5 μm, Cat. #773700-902) at a flow rate of 250 μL/min. Mobile phases A (2% acetonitrile in HPLC water) and B (98% acetonitrile in HPLC water) were used for RP gradient. MS analysis was performed on a Q-Exactive HF mass spectrometer (Thermo Fisher Scientific, USA) coupled to a Nanospray Flex source (Thermo Fisher Scientific, USA). The separated peptides were loaded onto a C18 column (50 cm × 75 μm) connected to an EASY-nLC™ 1200 system (Thermo Fisher Scientific, USA) and separated with a linear gradient of buffer B (80% acetonitrile and 0.1% formic acid) at a flow rate of 300 nL/min. The 1.5-h gradient was determined as follows: 8-25% buffer B for 70 min, 25-45% buffer B for 19 min, 45-100% buffer B for 6 min, and then 100% buffer B for a 10-min hold.

For the spectra library generation, the instrument was operated in DDA mode. MS scan was performed in the 350-1,650 m/z range at a resolution of 1.2×10^5^ with AGC target of 3×10^6^, and a maximum injection time of 100 ms, followed by selection of the 20 most abundant precursor ions for collisional induced dissociation (collision energy 27%) using a resolution of 3×10^4^ with an AGC target of 2×10^5^, a maximum injection time of 80 ms, and isolation window of 1.4 m/z. The Q-Exactive HF dynamic exclusion was set for 40.0 s and run under positive mode.

All tissue samples were then analyzed by DIA mode. MS scan was acquired in the mass range of 350-1250 m/z using a resolution of 1.2×10^4^ with AGC target of 3×10^6^, and a maximum injection time of 100 ms. Subsequently, the 32 acquisition windows were fragmented by higher energy collisional dissociation (collision energy 28%, each acquisition window of 26 m/z) using a resolution of 3 × 10^4^ with an AGC target of 1×10^6^, automatic maximum injection time, and run under positive mode.

### MS data analysis

Spectral libraries were generated using Spectronaut Pulsar software (Biognosys) by combining the output from DDA MS runs analyzed using Proteome Discoverer 2.0 (Thermo Scientific) and searched against Macaca_fascicularis_5.0 of the NCBI annotation. Spectronaut software was subsequently used to match raw DIA MS data against the spectral library using the default settings with slight modifications. The FDRs for protein, peptide, and peptide spectrum matches were all set to 0.01.

Protein expression values were normalized based on quantiles across all the samples for each tissue separately using normalize.quantiles in R package “preprocessCore”. Imputation was used to assign missing values to represent lack of detection based on a normal distribution. A simple linear model implemented in the R/Bioconductor package limma v3.42.2 ^76^ was used to identify the DEPs between diseased and healthy monkeys for each tissue separately. Proteins with fold change > 1.5 and Benjamini-Hochberg corrected p-values < 0.15 were considered to be differentially expressed.

### Pathway enrichment analysis and upstream regulator analysis

Ingenuity Pathway Analysis ^77^ content version 51963813 (Qiagen, Redwood City, USA) was used to identify over-represented pathways and upstream regulators for the DEGs and DEPs in each comparison. Canonical pathways with Benjamini-Hochberg corrected p-value < 0.05 and containing more than three DEGs were considered significantly overrepresented. Upstream regulators with Benjamini-Hochberg corrected p-value < 0.05 were considered significant.

### Molecular interaction network

NetworkAnalyst ^78^ was used to construct molecular networks for transcriptomics and proteomics data separately and also for two omics types combined. Lists of DEGs or/and DEPs were submitted to NetworkAnalyst to build a zero-order or a minimum-connected network, depending on the size of the lists. Tissue-specific protein-protein interactions were constructed using the DifferentialNet database if possible. Genes/proteins with a large number of molecular interactions were identified by counting the edges connecting the nodes. Pathway enrichment analysis was performed using all the genes/proteins in the network.

### Multi-omics integration analysis

To integrate across omics data and generate integrated transcriptomic and proteomic signature, we employed sparse generalized canonical correlation discriminant analysis via the DIABLO framework ^56^, which is implemented in the R package mixOmics version 6.14.1 ^79^. Normalized and log-transformed transcriptomic and proteomic data were used for the analysis. A value of 1 in the design matrix was determined using the “block.splsda” function. The optimal number of components and features included in the final model was determined by the “tune.block.splsda” function using the leave-one-out cross-validation. Relevance association networks were visualized in Cytoscape ^80^.

## Supplemental Information Titles and Legends

**Figure S1**. Transcriptomic analyses of obesity. (A) Tissue-specific transcriptomic profiles. Hierarchical clustering of the log transformed FPKQ values of all genes for monkeys in the non-obese group is shown in the heatmap. Expression values are scaled by row. Individual monkeys and genes are shown in columns and rows, respectively. (B-D) Volcano plots for DEGs in the abdominal WAT, BAT, and visceral WAT of obesity. The x-axis shows log2FCs between the obese and non-obese groups. Genes with increased and decreased expression in obesity are colored in green and pink, respectively. The y-axis shows the FDRs (Benjamini-Hochberg corrected p-values) on a -log10 scale. The horizontal dashed lines represent the cutoff of FDR < 0.05. The vertical dashed lines represent the cutoff of log2FC > log2(1.5) or < -log2(1.5). Genes with fold changes and p-values greater than the scale of the plot are plotted as triangles. The number of DEGs is shown in the upper corners of each plot. (E) The top 10 most significant pathways enriched in DEGs of the BAT in obesity. (F) Number of overlapping DEGs between tissues in obesity. Number of DEGs for each tissue are shown in the diagonal cells. Number of DEGs shared between tissues are shown in the other cells. BAT, brown adipose tissue; DEG, differentially expressed gene; FDR, false discovery rate; FPKQ, fragments per kilobase million quantile normalized; WAT, white adipose tissue.

**Figure S2**. Expression of *SLC2A1* (A) and *PCSK1* (B) across tissues in obesity. The x-axis shows the FPKQ of each sample, log2FC between the obese and non-obese monkeys, and -log10 transformed FDRs (Benjamini-Hochberg corrected p-values) respectively. Differential expression analysis was only performed for tissues with detectable expression (FPKQ > 1). The vertical dashed lines represent the cutoffs for DEGs: FPKQ > 1, FDR < 0.05, log2FC > log2(1.5) or < -log2(1.5). FDR, false discovery rate; FPKQ, fragments per kilobase million quantile normalized.

**Figure S3**. Proteomics analyses of obesity. (A) Number of confidence proteins in each tissue. (B) Hierarchical clustering of the proteomic data. (C-E) Volcano plots for DEPs in the abdominal WAT, BAT, and visceral WAT. The x-axis shows log2FCs between the obese and non-obese groups. Proteins with increased and decreased expression in obesity are colored in green and brown respectively. The y-axis shows the FDRs (Benjamini-Hochberg corrected p-values) on a -log10 scale. The horizontal dashed lines represent the cutoff of FDR < 0.15. The vertical dashed lines represent the cutoff of log2FC > log2(1.5) or < -log2(1.5). Proteins with fold changes and p-values greater than the scale of the plot are plotted as triangles. The number of DEPs is shown in the upper corners of each plot. (F) Venn diagram showing the number of overlapping DEPs between the adipose tissues for obesity. The total number of DEPs for each tissue is shown in the parentheses. (G) Protein-protein interaction network for DEPs of BAT of obesity. (H) Pathways significantly enriched in the protein-protein interaction network for the DEPs in the visceral WAT and BAT. The 10 most significant pathways for each tissue are shown. BAT, brown adipose tissue; DEP, differentially expressed protein; FDR, false discovery rate; WAT, white adipose tissue.

**Figure S4**. Multi-omics integrative analyses of obesity. (A) Radar plot showing the 27 shared DEGs and DEPs in the BAT of obesity. Error bars represent 95% confidence intervals. (B) Circos plot displaying all the DEGs and DEPs genome-wide in the BAT of obesity. On each chromosome, log2FCs of DEGs and DEPs and rainfall plots for DEGs and DEPs are vertically ordered from outside to inside tracks. (C) Heatmap of optimal discriminatory molecular signature for the BAT in obesity identified by the DIABLO method. (D) Relevance network constructed from the similarity matrix of the discriminatory molecular signature for the BAT to identify the association between proteins (blue circles) and RNA features (orange circles). Features exceeding a correlation coefficient cutoff of correlation coefficient (r) = 0.7 are considered associated and connected by lines. (E) Heatmap of optimal discriminatory molecular signature for the visceral WAT in obesity identified by the DIABLO method. (F) Circos plot showing the positive and negative correlation among the features in the discriminatory molecular signature of the visceral WAT based on a similarity matrix. Individual tiles on the ring represent individual genes. The connecting lines (orange or black) denote positive or negative correlations between genes with r > 0.7. BAT, brown adipose tissue; DEG, differentially expressed gene; DEP, differentially expressed protein; WAT, white adipose tissue.

**Figure S5**. Examples of an obesity-associated gene (A-C) and a T2D-associated gene (D). (A) UCSC Genome Browser snapshot showing the position of rs12254441 (yellow line) and its eQTL target gene *VDAC2* (pink box). (B-C) Contact heatmaps of Hi-C data showing the topological domains (TADs) for human pancreas and liver, respectively. Common variant rs12254441 is associated with BMI and mapped to gene *KAT6B* in the GWAS catalog. But it is an eQTL of *VDAC2* in the abdominal WAT in GTEx (TSS distance 352 kb) and is in the same topological domain with *VDAC2* in the liver and the pancreas, indicating *VDAC2* rather than *KAT6B* might be the target of rs12254441 and the gene associated with BMI. Hi-C data in (B) and (C) are from 3D Genome Browser (http://3dgenome.fsm.northwestern.edu/view.php). (D) Expression of *SLC2A2* across tissues in T2D. The x-axis shows the FPKQ of each sample, log2FC between the type 2 diabetic and non-diabetic (Non-T2D) monkeys, and -log10 transformed FDRs (Benjamini-Hochberg corrected p-values), respectively. *SLC2A2* was a DEG in the pancreas, liver, and renal medulla (in bold). Differential expression analysis was only performed for tissues with detectable expression (FPKQ > 1). The vertical dashed lines represent the cutoffs for DEGs: FPKQ >1, FDR < 0.01, log2FC > log2(1.5) or < -log2(1.5). BMI, body mass index; DEG, differentially expressed protein; FPKQ, fragments per kilobase million quantile normalized; T2D, type 2 diabetes; WAT, white adipose tissue.

**Figure S6**. Integrative analyses of obesity and T2D. (A) Relationships of WGCNA module eigengenes and clinical traits in the abdominal WAT. Co-expression modules identified by WGCNA are shown in rows and traits are in columns. Numbers in the cells are the correlations of the corresponding module eigengenes and traits. Cells are color coded by correlation according to the color legend. The raw p-values are shown in parentheses. The four modules significantly associated with disease state (either obesity or T2D) after Benjamini-Hochberg correction are shown in black boxes. BMI, body mass index; FBG, fasting blood glucose; HbA1c, glycated hemoglobin; AUC, area under the curve; TG, triglyceride; CHOL, total cholesterol. (B) Relationships of WGCNA module eigengenes and clinical traits in the visceral WAT. The two modules significantly associated with disease state (either obesity or T2D) after Benjamini-Hochberg correction are shown in black boxes. (C) Heatmap showing expression of the genes in the purple and darkred modules identified by WGCNA in the visceral WAT. Color bars indicate different modules and groups. Genes in the darkred module were downregulated in obesity and T2D but genes in the purple module were upregulated. (D) Significant pathways overrepresented in the genes of the purple module. (E) Significantly enriched pathways in the 62 shared DEPs showing same-direction changes between obesity and T2D in the abdominal WAT. T2D, type 2 diabetes; WAT, white adipose tissue; WGCNA, weighted-gene co-expression network analysis.

**Figure S7**. Transcriptional expression of *ISM1* and *SCL30A8*. (A) mRNA expression of *ISM1* in obesity. (B) mRNA expression of *ISM1* in T2D. (C) mRNA expression of *SLC30A8* in obesity. The x-axis shows the FPKQ of each sample, log2FC between the diseased and control monkeys, and -log10 transformed FDRs (Benjamini-Hochberg corrected p-values), respectively. Differential expression analysis was performed only for tissues with detectable expression (FPKQ > 1). The vertical dashed lines represent the cutoffs for DEGs: FPKQ >1, FDR < 0.05 in obesity and < 0.01 in T2D, log2FC > log2(1.5) or < -log2(1.5). DEG, differentially expressed protein; FDR, false discovery rate; FPKQ, fragments per kilobase million quantile normalized; T2D, type 2 diabetes.

**Table S1**. Summary of monkey cohorts collected in the present study.

**Table S2**. Expression datasets of human obesity in the public domain.

**Table S3**. GWAS DEGs in obesity.

**Table S4**. GWAS DEPs in obesity.

**Table S5**. Overlapping GWAS genes between DEGs and DEPs showing same-direction changes in T2D.

**Table S6**. GWAS DEGs of T2D identified in both GSE50244 and present study.

**Table S7**. Number of shared DEPs between obesity and T2D.

