## Supplemental Figures And Tables for "A Transcriptomic and Proteomic Atlas of Obesity and Type 2 Diabetes in Cynomolgus Monkeys"

**Figure S1**

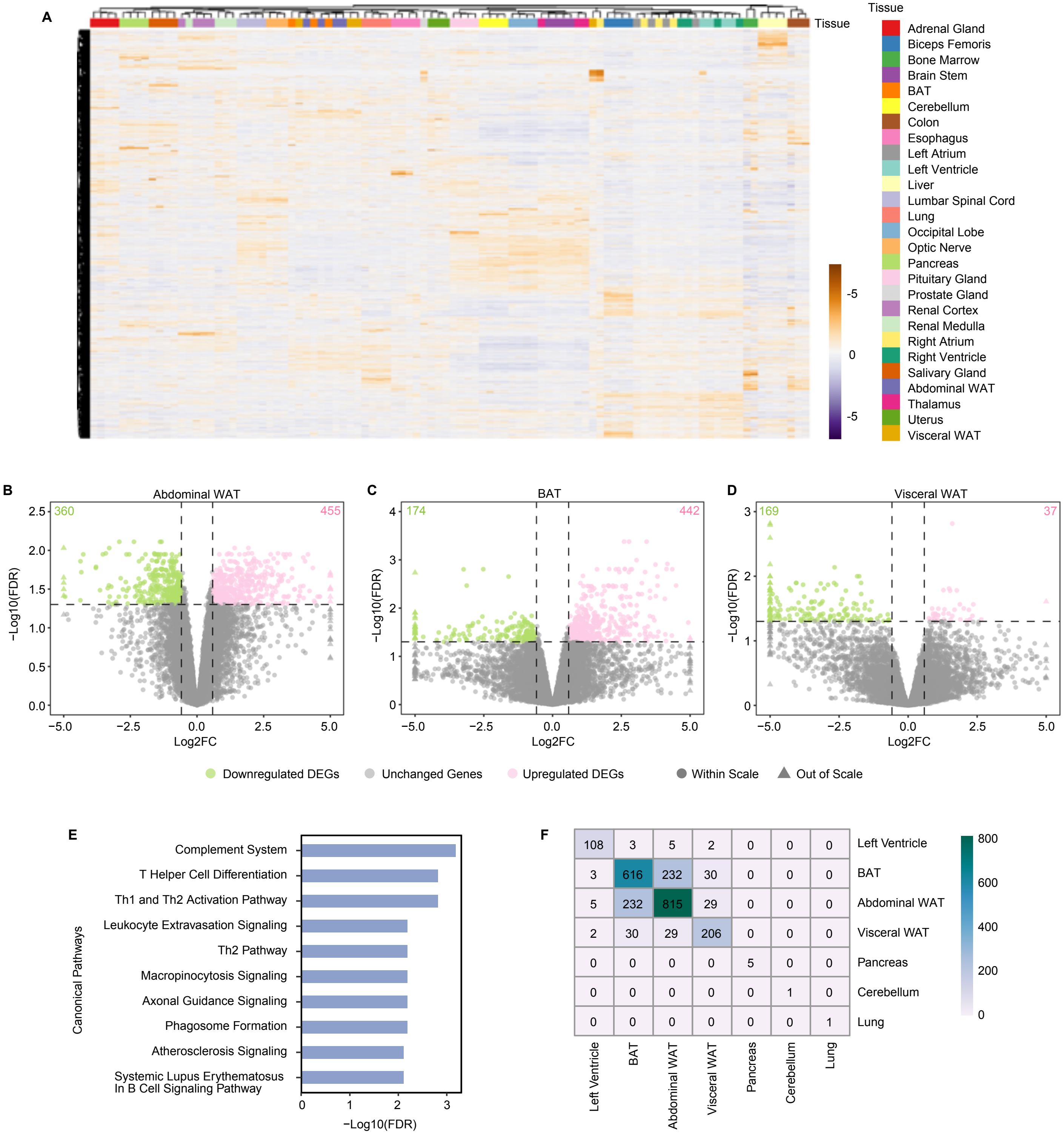
A
Tissue
Adrenal Gland
Biceps Femoris
Bone Marrow
Brain Stem
BAT
Cerebellum
Colon
Esophagus
Left Atrium
Left Ventricle
Liver
Lumbar Spinal Cord
Lung
Occipital Lobe
Optic Nerve
Pancreas
Pituitary Gland
Prostate Gland
Renal Cortex
Renal Medulla
Right Atrium
Right Ventricle
Salivary Gland
Abdominal WAT
Thalamus
Uterus
Visceral WAT
-5
0
-5
B
Abdominal WAT
360
455
C
BAT
174
442
D
Visceral WAT
169
37
−Log10(FDR)
Log2FC
Downregulated DEGs
Unchanged Genes
Upregulated DEGs
Within Scale
Out of Scale
E
Canonical Pathways
Complement System
T Helper Cell Differentiation
Th1 and Th2 Activation Pathway
Leukocyte Extravasation Signaling
Th2 Pathway
Macropinocytosis Signaling
Axonal Guidance Signaling
Phagosome Formation
Atherosclerosis Signaling
Systemic Lupus Erythematosus In B Cell Signaling Pathway
−Log10(FDR)
F
108 3 5 2 0 0 0 Left Ventricle
3 616 232 30 0 0 0 BAT
5 232 815 29 0 0 0 Abdominal WAT
2 30 29 206 0 0 0 Visceral WAT
0 0 0 0 5 0 0 Pancreas
0 0 0 0 0 1 0 Cerebellum
0 0 0 0 0 0 1 Lung
Left Ventricle
BAT
Abdominal WAT
Visceral WAT
Pancreas
Cerebellum
Lung
800
600
400
200
0

Figure S2

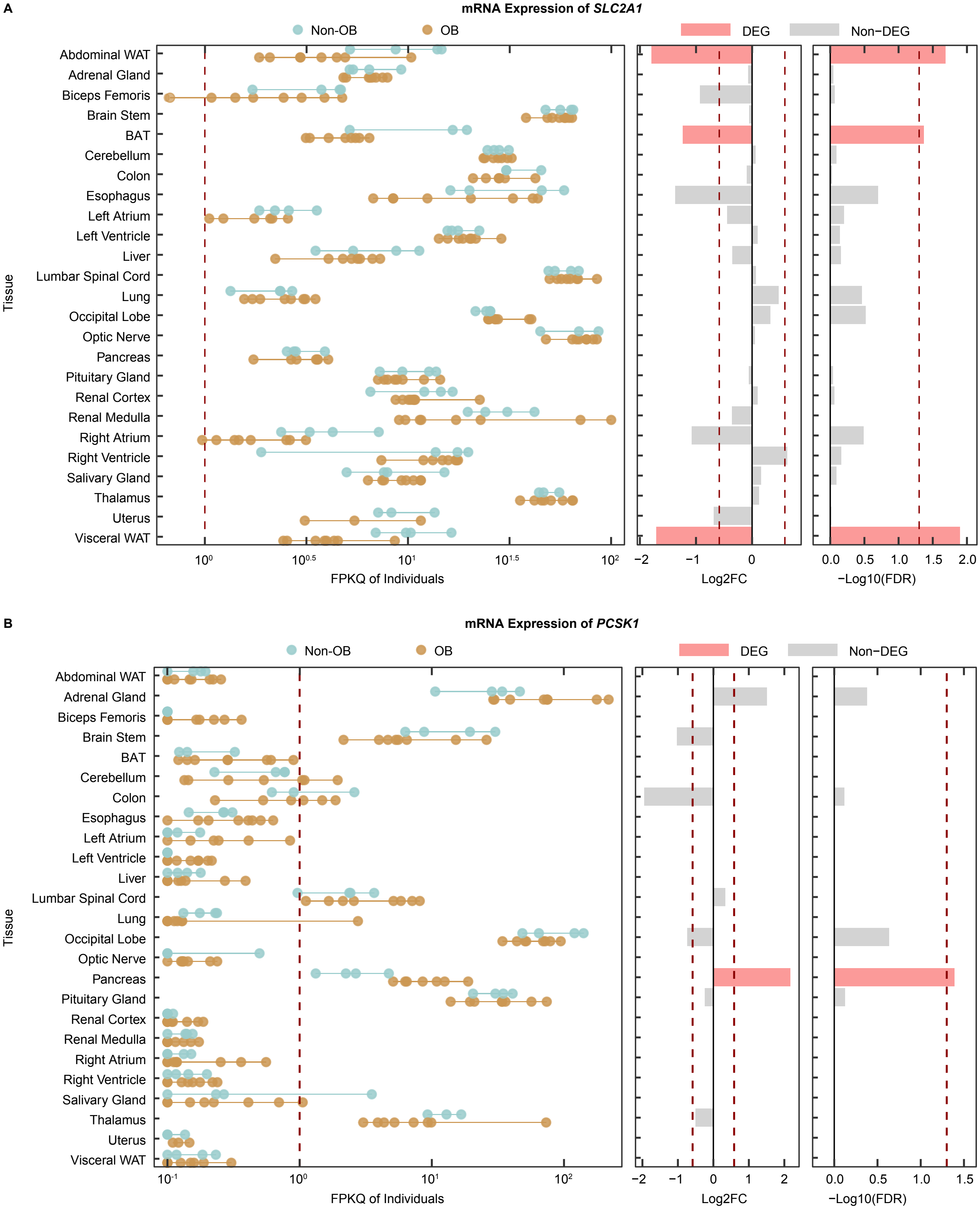
A
mRNA Expression of *SLC2A1*
Non-OB
OB
DEG
Non−DEG
Abdominal WAT
Adrenal Gland
Biceps Femoris
Brain Stem
BAT
Cerebellum
Colon
Esophagus
Left Atrium
Left Ventricle
Liver
Lumbar Spinal Cord
Lung
Occipital Lobe
Optic Nerve
Pancreas
Pituitary Gland
Renal Cortex
Renal Medulla
Right Atrium
Right Ventricle
Salivary Gland
Thalamus
Uterus
Visceral WAT
Tissue
$10^0$ $10^{0.5}$ $10^1$ $10^{1.5}$ $10^2$
FPKQ of Individuals
−2 −1 0 1
Log2FC
0.0 0.5 1.0 1.5 2.0
−Log10(FDR)
B
mRNA Expression of *PCSK1*
Non-OB
OB
DEG
Non−DEG
Abdominal WAT
Adrenal Gland
Biceps Femoris
Brain Stem
BAT
Cerebellum
Colon
Esophagus
Left Atrium
Left Ventricle
Liver
Lumbar Spinal Cord
Lung
Occipital Lobe
Optic Nerve
Pancreas
Pituitary Gland
Renal Cortex
Renal Medulla
Right Atrium
Right Ventricle
Salivary Gland
Thalamus
Uterus
Visceral WAT
Tissue
$10^{-1}$ $10^0$ $10^1$ $10^2$
FPKQ of Individuals
−2 −1 0 1 2
Log2FC
0.0 0.5 1.0 1.5
−Log10(FDR)

Figure S3

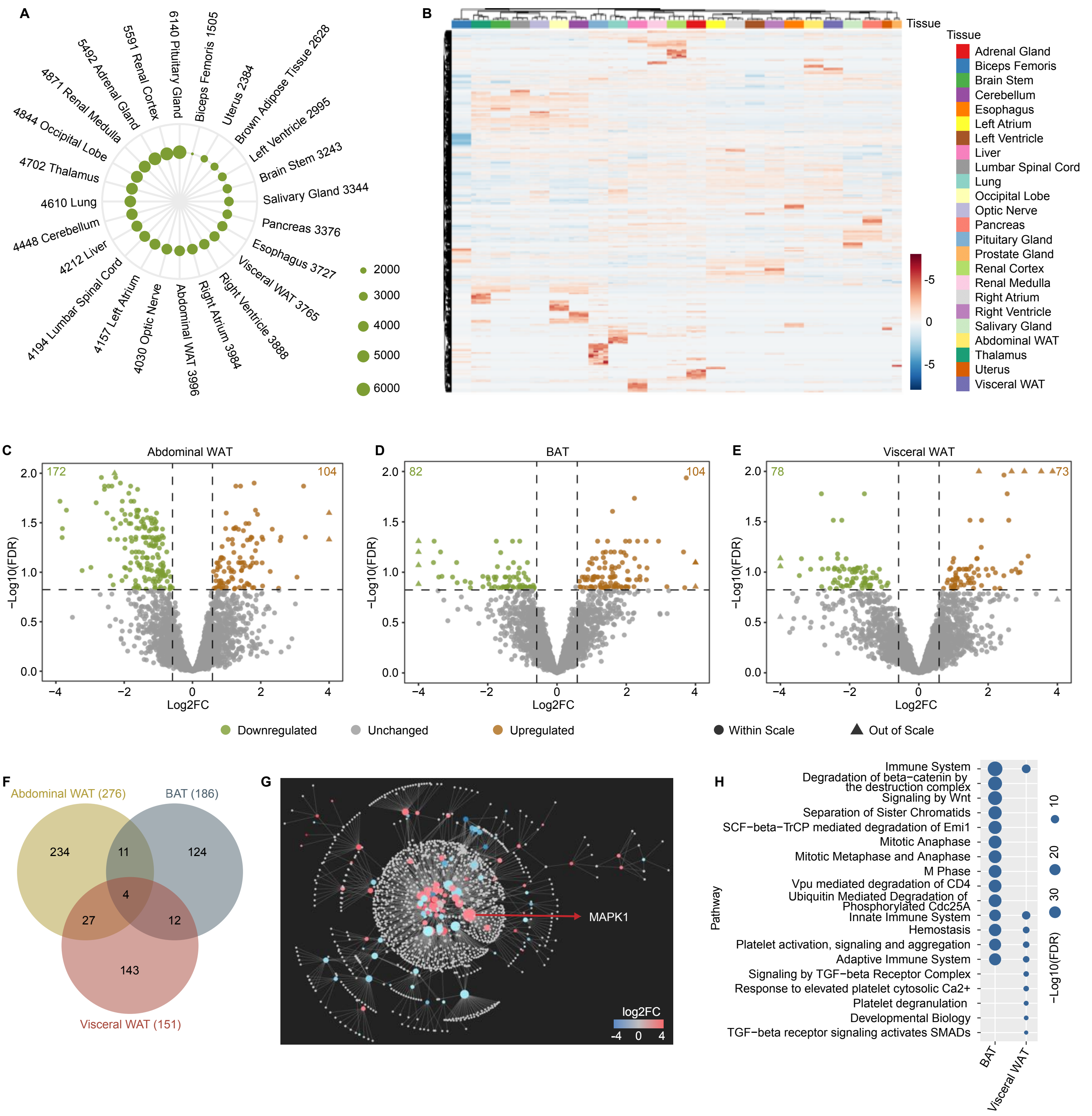
A
6140 Pituitary Gland
Biceps Femoris 1505
Uterus 2384
Brown Adipose Tissue 2628
Left Ventricle 2995
Brain Stem 3243
Salivary Gland 3344
Pancreas 3376
Esophagus 3727
Visceral WAT 3765
Right Ventricle 3888
Right Atrium 3984
Abdominal WAT 3996
4030 Optic Nerve
4157 Left Atrium
4194 Lumbar Spinal Cord
4212 Liver
4448 Cerebellum
4610 Lung
4702 Thalamus
4844 Occipital Lobe
4871 Renal Medulla
5492 Adrenal Gland
5591 Renal Cortex
2000
3000
4000
5000
6000
B
Tissue
Adrenal Gland
Biceps Femoris
Brain Stem
Cerebellum
Esophagus
Left Atrium
Left Ventricle
Liver
Lumbar Spinal Cord
Lung
Occipital Lobe
Optic Nerve
Pancreas
Pituitary Gland
Prostate Gland
Renal Cortex
Renal Medulla
Right Atrium
Right Ventricle
Salivary Gland
Abdominal WAT
Thalamus
Uterus
Visceral WAT
C
Abdominal WAT
172
104
D
BAT
82
104
E
Visceral WAT
78
73
−Log10(FDR)
Log2FC
Downregulated
Unchanged
Upregulated
Within Scale
Out of Scale
F
Abdominal WAT (276)
BAT (186)
Visceral WAT (151)
234
11
124
4
27
12
143
G
MAPK1
log2FC
-4 0 4
H
Immune System
Degradation of beta−catenin by the destruction complex
Signaling by Wnt
Separation of Sister Chromatids
SCF−beta−TrCP mediated degradation of Emi1
Mitotic Anaphase
Mitotic Metaphase and Anaphase
M Phase
Vpu mediated degradation of CD4
Ubiquitin Mediated Degradation of Phosphorylated Cdc25A
Innate Immune System
Hemostasis
Platelet activation, signaling and aggregation
Adaptive Immune System
Signaling by TGF−beta Receptor Complex
Response to elevated platelet cytosolic Ca2+
Platelet degranulation
Developmental Biology
TGF−beta receptor signaling activates SMADs
Pathway
BAT
Visceral WAT
−Log10(FDR)
10
20
30

**Figure S4**

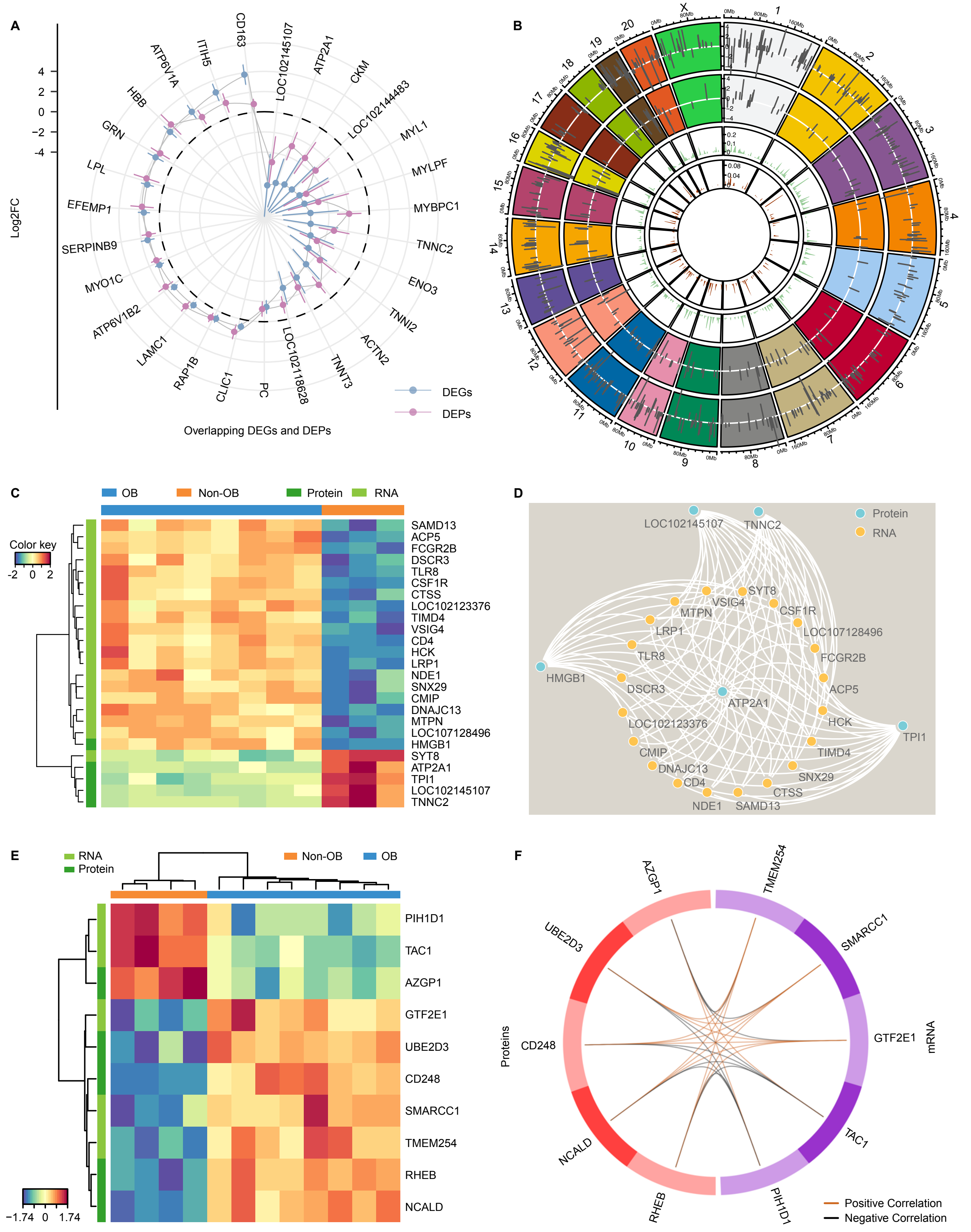
A
Log2FC
4
2
0
−2
-4
CD163
LOC102145107
ATP2A1
CKM
LOC102144483
MYL1
MYLPF
MYBPC1
TNNC2
ENO3
TNNI2
ACTN2
TNNT3
LOC102118628
PC
CLIC1
RAP1B
LAMC1
ATP6V1B2
MYO1C
SERPINB9
EFEMP1
LPL
GRN
HBB
ATP6V1A
ITIH5
DEGs
DEPs
Overlapping DEGs and DEPs
B
C
OB
Non-OB
Protein
RNA
Color key
-2
0
2
SAMD13
ACP5
FCGR2B
DSCR3
TLR8
CSF1R
CTSS
LOC102123376
TIMD4
VSIG4
CD4
HCK
LRP1
NDE1
SNX29
CMIP
DNAJC13
MTPN
LOC107128496
HMGB1
SYT8
ATP2A1
TPI1
LOC102145107
TNNC2
D
Protein
RNA
LOC102145107
TNNC2
SYT8
VSIG4
MTPN
CSF1R
LRP1
LOC107128496
TLR8
FCGR2B
HMGB1
DSCR3
ACP5
ATP2A1
HCK
LOC102123376
TPI1
CMIP
TIMD4
DNAJC13
SNX29
CD4
CTSS
NDE1
SAMD13
E
RNA
Protein
Non-OB
OB
PIH1D1
TAC1
AZGP1
GTF2E1
UBE2D3
CD248
SMARCC1
TMEM254
RHEB
NCALD
−1.74
0
1.74
F
AZGP1
TMEM254
UBE2D3
SMARCC1
CD248
GTF2E1
Proteins
mRNA
NCALD
TAC1
RHEB
PIH1D1
Positive Correlation
Negative Correlation

Figure S5

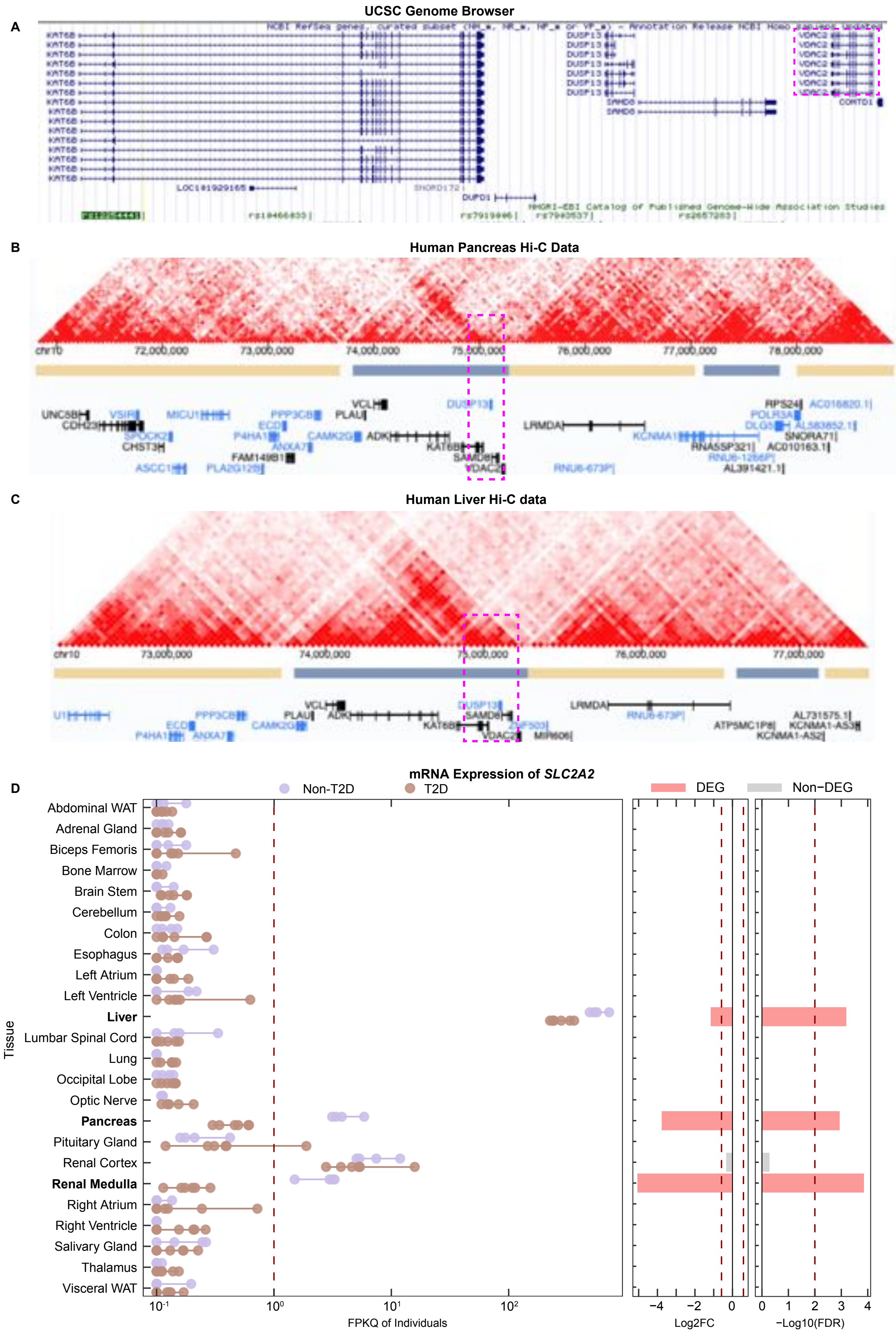
A
UCSC Genome Browser
KAT6B
DUSP13
VDAC2
SAMD8
COMTD1
LOC101929165
DUPD1
B
Human Pancreas Hi-C Data
chr10
72,000,000
73,000,000
74,000,000
75,000,000
76,000,000
77,000,000
78,000,000
UNC5B
VSIR
MICU1
PPP3CB
VCL
DUSP13
RPS24
AC016820.1
CDH23
PLAU
LRMDA
POLR3A
CHST3
ECD
ADK
KCNMA1
DLG5
AL583852.1
SPOCK2
P4HA1
CAMK2G
KAT6B
RNA5SP321
SNORA71
ANXA7
SAMD8
AC010163.1
FAM149B1
VDAC2
ASCC1
PLA2G12B
RNU6-673P
RNU6-1266P
AL391421.1
C
Human Liver Hi-C data
chr10
73,000,000
74,000,000
75,000,000
76,000,000
77,000,000
VCL
LRMDA
U1
PPP3CB
PLAU
ADK
DUSP13
RNU6-673P
AL731575.1
ECD
CAMK2G
KAT6B
SAMD8
ZNF503
ATP5MC1P8
KCNMA1-AS3
P4HA1
ANXA7
VDAC2
MIR606
KCNMA1-AS2
D
mRNA Expression of SLC2A2
Non-T2D
T2D
DEG
Non−DEG
Tissue
Abdominal WAT
Adrenal Gland
Biceps Femoris
Bone Marrow
Brain Stem
Cerebellum
Colon
Esophagus
Left Atrium
Left Ventricle
Liver
Lumbar Spinal Cord
Lung
Occipital Lobe
Optic Nerve
Pancreas
Pituitary Gland
Renal Cortex
Renal Medulla
Right Atrium
Right Ventricle
Salivary Gland
Thalamus
Visceral WAT
10⁻¹
10⁰
10¹
10²
FPKQ of Individuals
−4
−2
0
Log2FC
0
1
2
3
4
−Log10(FDR)

Figure S6

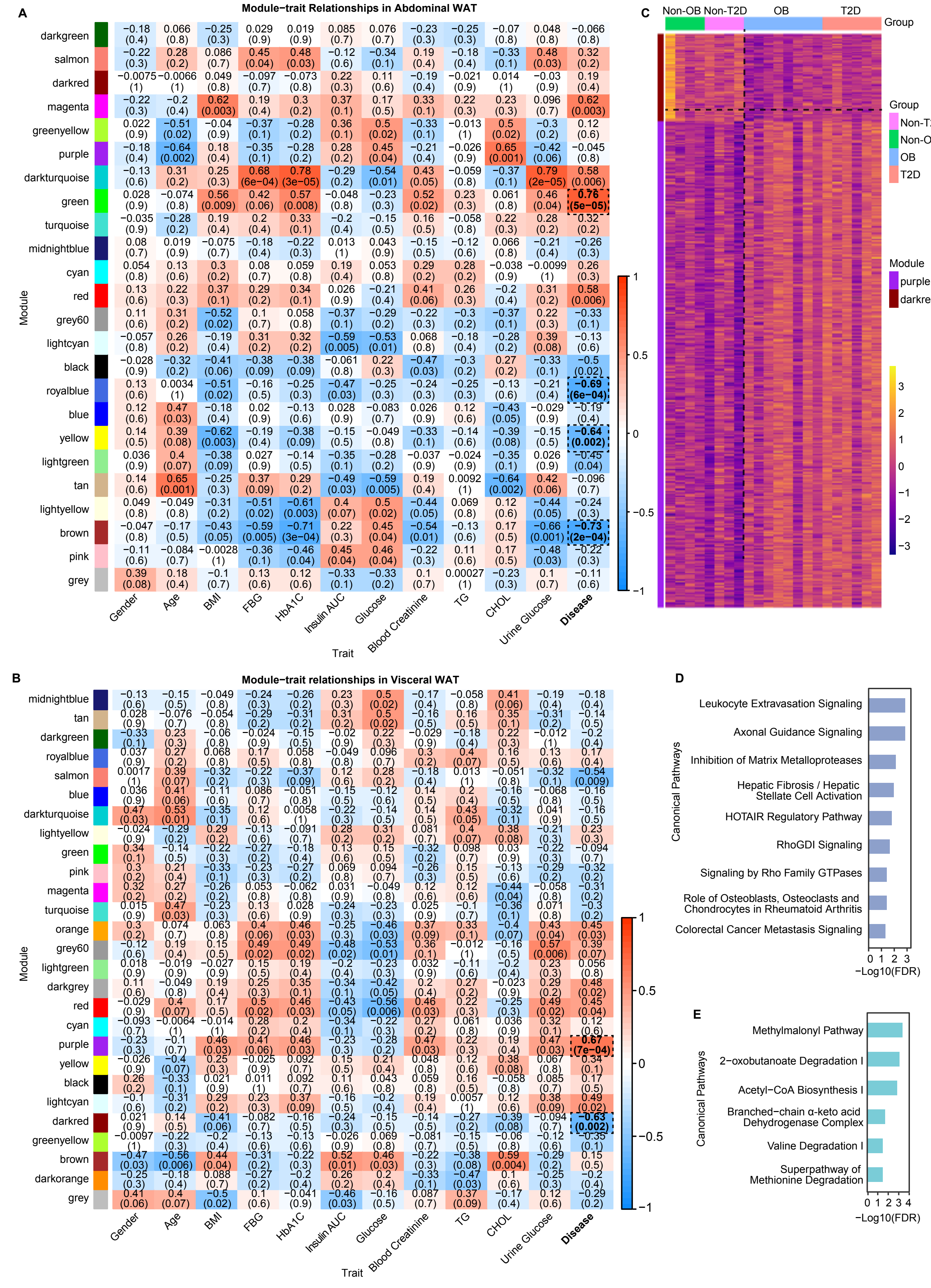
A
Module−trait Relationships in Abdominal WAT
darkgreen
salmon
darkred
magenta
greenyellow
purple
darkturquoise
green
turquoise
midnightblue
cyan
red
grey60
lightcyan
black
royalblue
blue
yellow
lightgreen
tan
lightyellow
brown
pink
grey
Module
Gender
Age
BMI
FBG
HbA1C
Insulin AUC
Glucose
Blood Creatinine
TG
CHOL
Urine Glucose
Disease
Trait
B
Module−trait relationships in Visceral WAT
midnightblue
tan
darkgreen
royalblue
salmon
blue
darkturquoise
lightyellow
green
pink
magenta
turquoise
orange
grey60
lightgreen
darkgrey
red
cyan
purple
yellow
black
lightcyan
darkred
greenyellow
brown
darkorange
grey
Module
Gender
Age
BMI
FBG
HbA1C
Insulin AUC
Glucose
Blood Creatinine
TG
CHOL
Urine Glucose
Disease
Trait
C
Non-OB
Non-T2D
OB
T2D
Group
Group
Non-T2D
Non-OB
OB
T2D
Module
purple
darkred
D
Leukocyte Extravasation Signaling
Axonal Guidance Signaling
Inhibition of Matrix Metalloproteases
Hepatic Fibrosis / Hepatic Stellate Cell Activation
HOTAIR Regulatory Pathway
RhoGDI Signaling
Signaling by Rho Family GTPases
Role of Osteoblasts, Osteoclasts and Chondrocytes in Rheumatoid Arthritis
Colorectal Cancer Metastasis Signaling
Canonical Pathways
−Log10(FDR)
E
Methylmalonyl Pathway
2−oxobutanoate Degradation I
Acetyl−CoA Biosynthesis I
Branched−chain α-keto acid Dehydrogenase Complex
Valine Degradation I
Superpathway of Methionine Degradation
Canonical Pathways
−Log10(FDR)

Figure S7

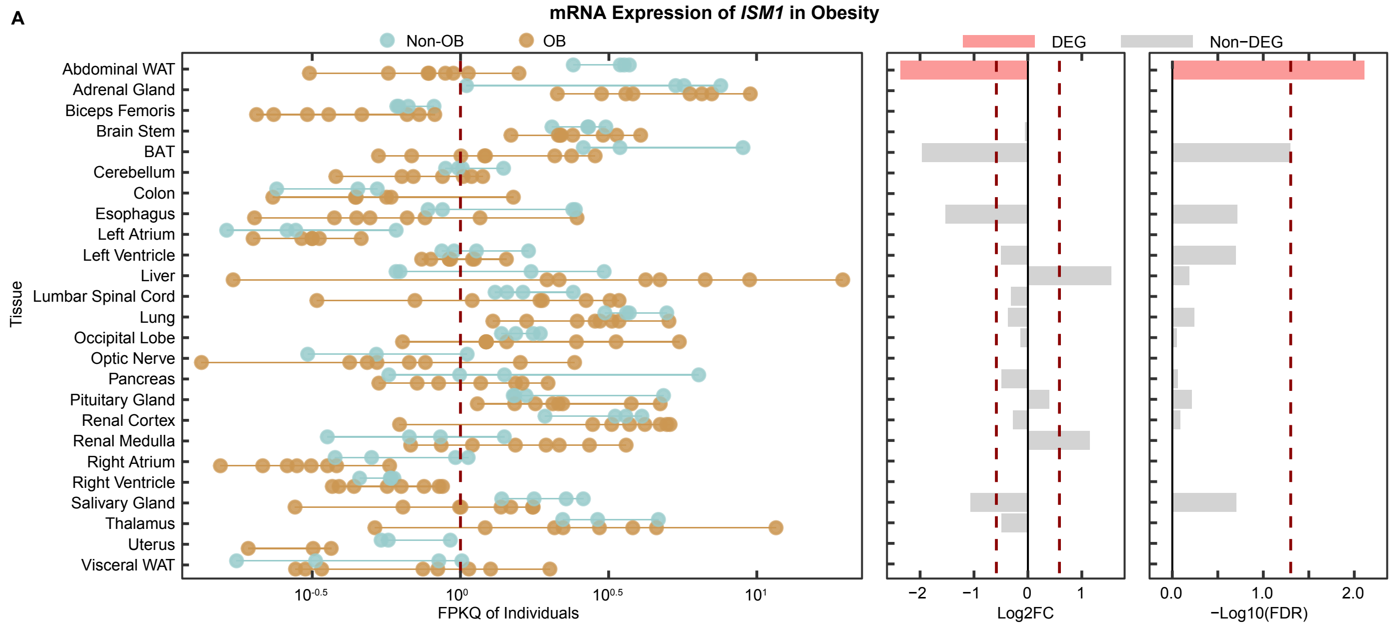

A
mRNA Expression of ISM1 in Obesity
Non-OB
OB
DEG
Non-DEG
Abdominal WAT
Adrenal Gland
Biceps Femoris
Brain Stem
BAT
Cerebellum
Colon
Esophagus
Left Atrium
Left Ventricle
Liver
Lumbar Spinal Cord
Lung
Occipital Lobe
Optic Nerve
Pancreas
Pituitary Gland
Renal Cortex
Renal Medulla
Right Atrium
Right Ventricle
Salivary Gland
Thalamus
Uterus
Visceral WAT
Tissue
FPKQ of Individuals
Log2FC
−Log10(FDR)


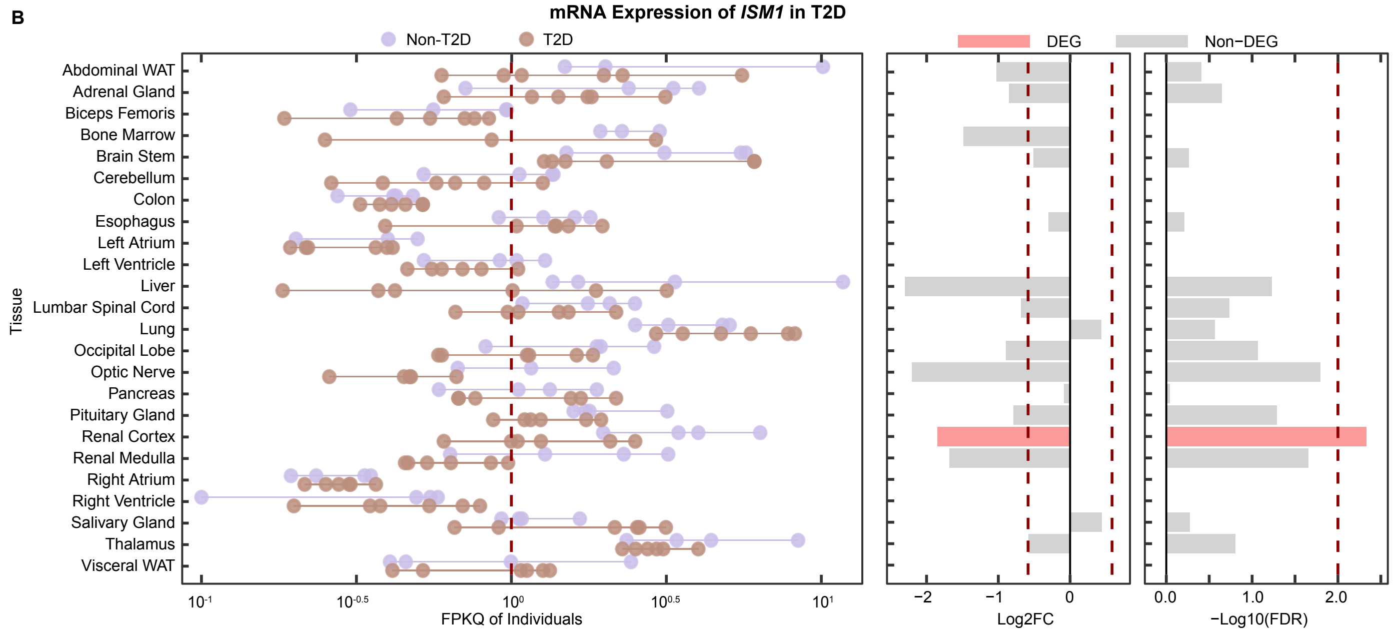

B
mRNA Expression of ISM1 in T2D
Non-T2D
T2D
DEG
Non-DEG
Abdominal WAT
Adrenal Gland
Biceps Femoris
Bone Marrow
Brain Stem
Cerebellum
Colon
Esophagus
Left Atrium
Left Ventricle
Liver
Lumbar Spinal Cord
Lung
Occipital Lobe
Optic Nerve
Pancreas
Pituitary Gland
Renal Cortex
Renal Medulla
Right Atrium
Right Ventricle
Salivary Gland
Thalamus
Visceral WAT
Tissue
FPKQ of Individuals
Log2FC
−Log10(FDR)


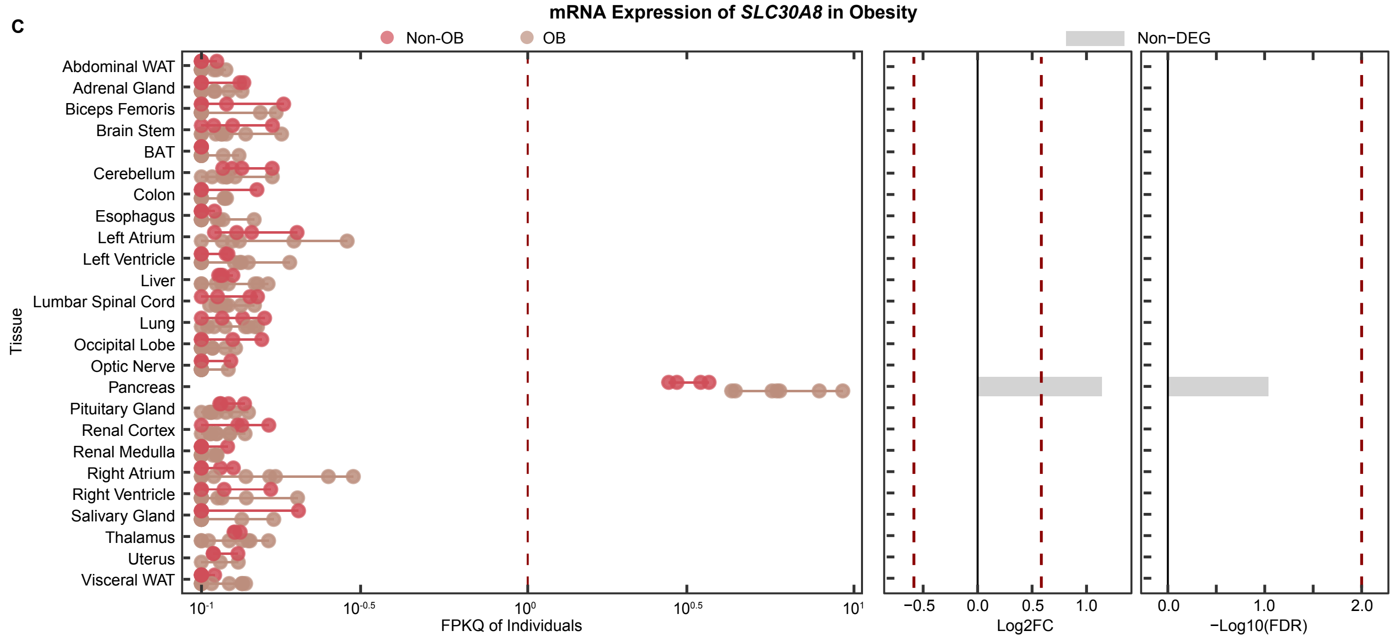

C
mRNA Expression of SLC30A8 in Obesity
Non-OB
OB
Non-DEG
Abdominal WAT
Adrenal Gland
Biceps Femoris
Brain Stem
BAT
Cerebellum
Colon
Esophagus
Left Atrium
Left Ventricle
Liver
Lumbar Spinal Cord
Lung
Occipital Lobe
Optic Nerve
Pancreas
Pituitary Gland
Renal Cortex
Renal Medulla
Right Atrium
Right Ventricle
Salivary Gland
Thalamus
Uterus
Visceral WAT
Tissue
FPKQ of Individuals
Log2FC
−Log10(FDR)

Table S1. Summary of monkey cohorts collected in the present study.

| Group | Animal ID | Gender | Age | BMI (kg/m2) | FBG (mg/dL) | HbA1C (%) | Triglycerides (mM/L) | CHOL (mM/L) | Blood Creatinine (μM/L) | Urine Glucose |
|---|---|---|---|---|---|---|---|---|---|---|
| **Non-OB** | YC1 | F | 13 | 21.54 | 56.16 | 5.07 | 0.75 | 2.73 | 89 | 0 |
| | YC2 | F | 11 | 22.68 | 79.38 | 4.97 | 0.48 | 3.97 | 61 | 0 |
| | YC3 | F | 11 | 17.55 | 53.10 | 4.81 | 0.67 | 3.69 | 59 | 0 |
| | OW2 | M | 8 | 28.20 | 62.64 | 5.41 | 0.25 | 1.66 | 54 | 0 |
| | **Mean ± SD** | | **10.8±2.1** | **22.5±4.4** | **62.8±11.7** | **5.1±0.3** | **0.54±0.22** | **3.0±1.1** | **65.8±15.8** | **0** |
| **OB** | OB1 | M | 6 | 47.97 | 47.88 | 5.59 | 0.52 | 1.9 | 116 | 0 |
| | OB2 | M | 8 | 43.19 | 51.84 | 5.42 | 0.65 | 2.93 | 101 | 0 |
| | OB3 | F | 12 | 35.63 | 50.40 | 5.31 | 1.37 | 3.03 | 72 | 0 |
| | OW1 | M | 7 | 31.30 | 51.30 | 5.32 | 0.59 | 3.51 | 111 | 0 |
| | PD1 | M | 7 | 36.36 | 48.24 | 5.70 | 0.34 | 2.31 | 121 | 0 |
| | PD2 | F | 10 | 40.78 | 54.18 | 6.16 | 1.11 | 3.86 | 48 | 0 |
| | PD3 | F | 9 | 37.06 | 57.42 | 5.86 | 3.24 | 3.66 | 81 | 0 |
| | PD4 | M | 7 | 33.01 | 72.18 | 5.74 | 0.49 | 2.05 | 97 | 0 |
| | **Mean ± SD** | | **8.3±2.0** | ***38.2±5.5*** | **54.2±7.9** | ***5.6±0.3**** | **1.04±0.95** | **2.9±0.8** | **93.4±24.9** | **0** |
| **Non-T2D** | AC1 | M | 16 | 28.99 | 89.10 | 4.90 | 0.51 | 2.68 | 57 | 0 |
| | AC2 | M | 18 | 28.44 | 67.68 | 4.30 | 0.42 | 2.12 | 99.2 | 0 |
| | AC3 | F | 17 | 17.23 | 69.30 | 4.30 | 0.37 | 3.38 | 44.8 | 0 |
| | AC4 | F | 15 | 15.97 | 58.32 | 4.30 | 0.26 | 2.16 | 37.2 | 0 |
| | **Mean ± SD** | | **16.5±1.3** | **22.7±7.0** | **71.1±12.9** | **4.5±0.3** | **0.39±0.10** | **2.6±0.6** | **59.6±27.7** | **0** |
| **T2D** | DM1 | M | 16 | 51.99 | 342.00 | 11.80 | 11.13 | 4.51 | 56.6 | 4 |
| | DM2 | M | 14 | 40.97 | 217.62 | 11.70 | 1.6 | 2.11 | 46.9 | 4 |
| | DM3 | M | 17 | 34.32 | 287.46 | 12.10 | 4.93 | 4.88 | 34.6 | 4 |
| | DM4 | F | 18 | 24.85 | 252.36 | 12.60 | 1.87 | 2.66 | 42.4 | 4 |
| | DM5 | F | 16 | 24.19 | 244.80 | 10.50 | 2.87 | 2.02 | 42.4 | 4 |
| | DM6 | F | 16 | 22.81 | 140.94 | >14.0 | 0.49 | 2.63 | 26 | 4 |
| | **Mean ± SD** | | **16.2±1.3** | **33.2±11.6** | ***248±68**** | ***12.1±1.2**** | ***3.82±3.88**** | **3.1±1.2** | **41.5±10.5** | ***4**** |

BMI, Body Mass Index; FBG, Fasting Blood Glucose; HbA1C, Glycated Hemoglobin; CHOL, Total Cholesterol.
Non-OB, Non-obesity; OB, Obesity; T2D, Type 2 Diabetes; Non-T2D, Non-Type 2 Diabetes.
Statistical significance is denoted as * p-value < 0.05 and ** p-value < 0.01 by Mann-Whitney U test between diseased groups and non-diseased groups.

Table S2. Expression datasets of human obesity in the public domain.

| GEO | Tissue | Sample | Data Type | Journal | Year | Number of DEGs (FDR < 0.05) | Number of DEGs (FDR < 0.05 & FC > 1.2) |
|---|---|---|---|---|---|---|---|
| GSE2508 | Abdominal subcutaneous adipocytes | 19 cases vs. 20 controls | Array | Diabetologia | 2005 | 2412 | 2329 |
| GSE15653 | Liver | 4 cases vs. 5 controls | Array | J Clin Endocrinol Metab | 2009 | 1 | 0 |
| GSE29718 | Subcutaneous white adipose tissue | 7 cases vs. 8 controls | Array | Int J Pediatr Obes | 2011 | 0 | 0 |
| GSE48964 | Subcutaneous white adipose tissue | 3 cases vs. 3 controls | Array | BMC Genomics | 2013 | 0 | 0 |
| GSE48452 | Liver | 27 cases vs. 14 controls | Array | Cell Metab | 2013 | 2 | 0 |
| GSE9624 | Visceral adipose tissue | 14 cases vs. 13 controls | Array | Int J Mol Sci | 2015 | 0 | 0 |
| GSE81965 | Primary differentiated myotubes from skeletal muscle | 6 cases vs. 6 controls | RNA-seq | Genome Medicine | 2017 | 5145 | 184 |
| Not available | Breast tissue | 31 cases vs. 31 controls | RNA-seq | Breast Cancer Research | 2020 | 120 | 120 |
| GSE155960 | Stromal vascular fraction of human white adipose tissue | 3 cases vs. 3 controls | scRNA-seq | Nature Immunology | 2021 | 15-740 | |

Table S3. GWAS DEGs in obesity.

| Gene | Chr | Start | End | Tissue | Change in Obesity |
|---|---|---|---|---|---|
| *ASAH1* | 8 | 18,078,401 | 18,108,329 | Abdominal WAT & BAT | Upregulated |
| *CMKLR1* | 11 | 111,926,335 | 111,979,484 | Abdominal WAT & BAT | Upregulated |
| *HOXB3* | 16 | 33,902,522 | 33,937,441 | Abdominal WAT & BAT | Downregulated |
| *ECM1* | 1 | 101,559,204 | 101,565,359 | BAT | Upregulated |
| *KIAA1211* | 5 | 78,087,204 | 78,378,919 | BAT | Downregulated |
| *KCNMA1* | 9 | 57,985,995 | 58,749,040 | BAT | Downregulated |
| *ADCY3* | 13 | 86,001,639 | 86,117,392 | BAT | Downregulated |
| *STAC* | 2 | 10,858,910 | 11,031,792 | Abdominal WAT | Downregulated |
| *P2RY1* | 2 | 55,214,310 | 55,219,470 | Abdominal WAT | Upregulated |
| *CARD11* | 3 | 37,917,677 | 38,057,108 | Abdominal WAT | Upregulated |
| *BCKDHB* | 4 | 93,506,906 | 93,751,455 | Abdominal WAT | Downregulated |
| *ITPR3* | 4 | 137,119,373 | 137,194,853 | Abdominal WAT | Downregulated |
| *ZDBF2* | 12 | 95,869,866 | 95,909,328 | Abdominal WAT | Downregulated |
| *IRS1* | 12 | 116,526,101 | 116,589,153 | Abdominal WAT | Downregulated |
| *LRRFIP1* | 12 | 127,453,108 | 127,622,688 | Abdominal WAT | Upregulated |
| *PTDSS2* | 14 | 264,657 | 306,582 | Abdominal WAT | Downregulated |
| *CMYA5* | 6 | 77,987,969 | 78,092,127 | Visceral WAT | Downregulated |
| *FAIM2* | 11 | 48,726,441 | 48,757,478 | Visceral WAT | Downregulated |
| *SLC8A1* | 13 | 70,016,851 | 70,420,351 | Visceral WAT | Downregulated |
| *PRL* | 4 | 148,342,109 | 148,357,466 | Cerebellum | Downregulated |
| *DNAJC27* | 13 | 85,945,379 | 85,975,102 | Left Ventricle | Downregulated |

Shown are the overlapping genes between the DEGs obtained from the differential expression analysis between obesity and young control groups and the obesity-associated genes in the GWAS catalog. DEG, differentially expressed gene.

Table S4. GWAS DEPs in obesity.

| Gene | Chr | Start | End | Tissue | Change in Obesity |
|---|---|---|---|---|---|
| C7 | 6 | 41,396,080 | 41,478,907 | Visceral WAT | Downregulated |
| CFB | 4 | 138,901,319 | 138,907,529 | Visceral WAT | Downregulated |
| CFH | 1 | 54,926,945 | 55,183,655 | BAT | Upregulated |
| ECM1 | 1 | 101,559,204 | 101,565,359 | Abdominal WAT | Upregulated |
| FHL1 | X | 132,912,255 | 132,983,580 | Abdominal WAT | Upregulated |
| PTER | 9 | 16,983,737 | 17,058,693 | Uterus | Upregulated |
| RTN4 | 13 | 55,009,654 | 55,090,659 | BAT & Visceral WAT | Upregulated |
| TGM2 | 10 | 69,712,824 | 69,751,781 | Visceral WAT | Downregulated |

Shown are the DEPs obtained from the differential protein expression analysis between obesity and young control groups and associated with obesity in the GWAS catalog.

Table S5. Overlapping GWAS genes between DEGs and DEPs showing same-direction changes in T2D.

| **Tissue** | **Number of Genes** | **Gene Name** |
|---|---|---|
| Renal Cortex | 19 | *ANPEP, BOP1, C1GALT1, CADM1, COG7, FMO4, GNPDA2, HMGA1, LGALSL, MRPL13, MTAP, PKLR, PRKD1, PVRL2, SLC25A51, TBC1D4, TOMM40, UBLCP1, VTA1* |
| Renal Medulla | 17 | *ANPEP, AOC1, CDH2, DMGDH, HMGB1, KDM4B, LGALSL, LPP, MBNL1, PABPC4, PSMD12, SLCO4C1, SUGCT, TIMM17B, TINAGL1, TMEM167A, TTN* |
| Left Atrium | 16 | *ARL15, DNAJC2, FCGRT, GNPDA2, IMPA2, LGALSL, MOB1B, MRPS30, MTAP, PDE3A, PLCB3, QKI, SCN7A, SGCD, TRIM63, TRIOBP* |
| Cerebellum | 12 | *CADM2, G6PD, GPSM1, KATNAL1, LAMB2, MACROD2, MRPL13, MTAP, SCYL1, TOMM40, WFS1, YTHDF3* |
| Left Ventricle | 9 | *APIP, COPB1, GNAI2, HMGB1, LPL, MRPL13, MRPL19, OLA1, RPL13* |
| Occipital Lobe | 9 | *DAAM1, DGKB, MDGA2, MRPL13, MYO1C, PITPNM2, RBM17, SCYL1, ZC3H4* |
| Liver | 7 | *APOE, DNAJC2, HAT1, LAMB2, LHPP, QKI, UBLCP1* |
| Right Ventricle | 7 | *CASKIN2, GNPDA2, HSD17B12, LPL, MRPL13, PSMC2, TOM1* |
| Right Atrium | 6 | *ATP2A3, COPB1, GNPDA2, PDE3A, SCN7A, TINAGL1* |
| Salivary Gland | 6 | *COPB1, HMGA1, HSD17B12, MGAT1, PEPD, STARD10* |
| Lumbar Spinal Cord | 4 | *HSD17B12, KDM4B, PEPD, TMEM106B* |
| Pituitary Gland | 4 | *CCDC92, HPCAL1, PIK3R1, RPTOR* |
| Brain Stem | 1 | *TOMM40* |
| Optic Nerve | 1 | *DOCK4* |

Shown are the overlapping genes between the DEPs and DEGs showing same-direction changes in T2D and also associated with T2D in the GWAS catalog.

Table S6. GWAS DEGs of T2D identified in both GSE50244 and present study.

| Gene | Chr | Start | End | Log2 Fold Change (T2D/Control) | p-value | Adjusted p-value |
|---|---|---|---|---|---|---|
| *SLC2A2* | 2 | 75,192,208 | 75,224,946 | -3.76 | 3.19E-07 | 1.16E-03 |
| *ST18* | 8 | 53,070,682 | 53,398,358 | -3.22 | 8.70E-05 | 5.48E-03 |
| *HSF1* | 8 | 145,943,775 | 145,970,916 | 1.18 | 1.78E-05 | 3.61E-03 |
| *WSCD2* | 11 | 111,760,376 | 111,889,838 | -2.40 | 3.99E-05 | 4.42E-03 |
| *RAMP1* | 12 | 127,693,700 | 127,751,234 | 1.91 | 9.98E-05 | 5.65E-03 |
| *HPCAL1* | 13 | 100,548,812 | 100,669,923 | 0.93 | 3.28E-04 | 8.85E-03 |
| *MAP3K11* | 14 | 8,870,699 | 8,888,660 | 1.33 | 3.46E-06 | 2.39E-03 |
| *LLGL2* | 16 | 72,822,353 | 72,873,778 | 1.22 | 4.16E-06 | 2.51E-03 |
| *TCF3* | 19 | 1,419,686 | 1,473,680 | 1.11 | 2.70E-04 | 8.16E-03 |
| *RPL13* | 20 | 78,001,573 | 78,004,068 | 0.98 | 4.46E-04 | 9.90E-03 |

Table S7. Number of shared DEPs between obesity and T2D.

| Tissue | Number of Shared DEPs | Number of Shared DEPs Showing Same-direction change | Number of Shared DEPs Showing Opposite-direction change |
|---|---|---|---|
| Abdominal WAT | 71 | 62 | 9 |
| Visceral WAT | 21 | 7 | 14 |
| Cerebellum | 10 | 4 | 6 |
| Occipital Lobe | 3 | 3 | 0 |
| Pituitary Gland | 2 | 0 | 2 |
| Lung | 1 | 1 | 0 |
| Right Atrium | 1 | 0 | 1 |
